# DAF-18 is required for the age-dependent increase in DAF-16 activity in *Caenorhabditis elegans*

**DOI:** 10.1101/2020.11.16.384529

**Authors:** Kali Carrasco, Matthew J. Youngman

## Abstract

The insulin/insulin-like growth factor signaling (IIS) pathway modulates growth, survival, and lifespan by regulating FOXO transcription factors. In *Caenorhabditis elegans*, IIS maintains DAF-16/FOXO in an inactive state unless animals are challenged by environmental stress. Recent evidence suggests that DAF-16 becomes activated as part of normal aging in *C. elegans*, yet the regulatory module responsible for this phenomenon is largely undefined. Embedded within IIS is phospholipid signaling in which PIP_3_ produced by the PI3 kinase AGE-1 is an upstream event in DAF-16 inhibition. Countering AGE-1 is DAF-18, an ortholog of human PTEN phosphatase that dephosphorylates PIP_3_. Although it is required for normal lifespan in *C. elegans*, functional characterization of DAF-18 has primarily focused on its roles during development in the germline and neurons. In this study we asked whether DAF-18 plays a role in the age-dependent activation of DAF-16, and specifically in DAF-16-mediated immunity. Our data show that DAF-18 is expressed in multiple tissues during adulthood. We found that DAF-18 contributes to host defense in adult animals by functioning in the neurons and intestine, likely working through DAF-16 which acts in those same tissues to confer immunity. Supporting this possibility, DAF-18 was required for increased DAF-16 transcriptional activity during aging. Post-translational modifications including ubiquitination and sumoylation appear to be required for the function of DAF-18 during aging in *C. elegans*, indicating that strategies to modulate PTEN activity are evolutionarily conserved. Our results establish an important role for DAF-18 later in life and imply that it is a critical component of a neuroendocrine signaling circuit that governs the dynamic activity of DAF-16.

## INTRODUCTION

In diverse animal species, the insulin and insulin-like growth factor signaling pathway (IIS) regulates lifespan and integrates environmental cues to control growth and development (Kenyon *et al*. 1993; Tamemoto *et al*. 1994; Kimura *et al*. 1997; Tatar *et al*. 2001; Yuan *et al*. 2009). While eliminating insulin signaling reduces the size of some organisms, lifespan extension is a benefit in common to animals with low IIS activity. This was demonstrated first in the roundworm *Caenorhabditis elegans* but has since been shown to hold true in vertebrates as well (Kenyon *et al*. 1993). For example, female mice heterozygous for mutations in the insulin-like growth factor receptor *IGF1R* live 33% longer than their wildtype counterparts (Holzenberger *et al*. 2003). In dogs, a nonsynonymous SNP in *IGF1R* that may reduce the ability of the receptor to bind its ligand is found in tiny breeds, which tend to live longer, and is not present in large breeds that die comparatively younger (Hoopes *et al*. 2012). In humans, mutations in *IGF1R* associated with reduced IGF1R activity are found in centenarians (Suh *et al*. 2008). Regulatory mechanisms that attenuate insulin signaling could therefore be of particular importance to preserving normal lifespan.

In *C. elegans*, the basis for the effect of IIS on influencing longevity is its regulation of the ultimate downstream target of the pathway, the forkhead box class O (FOXO) transcription factor DAF-16. When bound by agonist insulin-like peptides, the kinase activity of the insulin receptor DAF-2 is activated and it phosphorylates the PI3 kinase AGE-1. AGE-1 phosphorylates PIP_2_ to produce PIP_3_, which is necessary to activate PDK-1, which, in turn, activates the kinase AKT-1. AKT-1 is the component of the IIS pathway that directly interacts with and inhibits DAF-16. When phosphorylated by AKT-1, DAF-16 is expelled from the nucleus and sequestered in the cytosol, physically separated from its transcriptional targets. Mutations in *daf-2* or *age-1* that disrupt the upstream steps of the IIS pathway extend the lifespan of *C. elegans* by 2- to 3-fold (Kenyon *et al*. 1993; Dorman *et al*. 1995). This extraordinary phenotype is suppressed by loss-of-function mutations in *daf-16*, indicating that its activity is what is responsible for augmenting the longevity of *daf-2* and *age-1* mutants (Kenyon *et al*. 1993). The transcriptional targets of DAF-16 are well-defined, and they span broad functional categories including metabolism, detoxification, genome maintenance, proteostasis, and innate immunity (McElwee *et al*. 2003; Murphy *et al*. 2003; Tepper *et al*. 2013; Li *et al*. 2019). Many DAF-16 targets play important roles in the response to environmental insults, and so activation of DAF-16 is correlated with resistance to ultraviolet light, oxidative stress, and microbial infection (Henderson and Johnson 2001; Wolff *et al*. 2006). By upregulating repair and maintenance pathways, DAF-16 improves cellular health, limits cellular damage, buffers organisms from stress and thus increases healthspan—the period of life spent in youthful vigor and without disease.

When wildtype *C. elegans* are maintained under stress-free conditions and their food source is unlimited, constitutive inhibition through IIS prevents DAF-16 from being activated. Until recently, environmental insults were the only inputs known to shift DAF-16 into an active state. Three reports now challenge that notion by suggesting that aging is sufficient to upregulate DAF-16 expression and to induce its transcriptional activity (Bansal *et al*. 2014; Li *et al*. 2019; McHugh *et al*. 2020). The regulatory mechanism that governs the dynamic activity of DAF-16 during aging is, at this point, largely undefined.

The series of successive activation events in the IIS pathway that lead to DAF-16 inhibition is reminiscent of similar modules in other signaling pathways that are subject to redundant regulation. One example is the familiar phosphorylation cascade within MAPK signaling pathways. Kinases within these pathways may be inactivated through dephosphorylation by members of three different classes of protein phosphatases. For example, ERK1/2 is the sole substrate of three dual-specificity phosphatases of the MKP family, but it is also regulated by an additional six phosphatases that dephosphorylate it and at least one other type of MAPK (Kondoh and Nishida 2007). There are considerably fewer phosphatases that act to inhibit IIS components. These proteins are attractive candidates for putative regulators of the age-dependent activation of DAF-16. Perhaps because of its broad spectrum of substrates, AKT-1 is the most tightly regulated IIS kinase, being prone to dephosphorylation by two tyrosine phosphatases, EAK-6 and SDF-9, and one serine/threonine kinase (of the PP2A family) (Hu *et al*. 2006; Padmanabhan *et al*. 2009). In contrast, the PI3 kinase AGE-1 is regulated by only a single phosphatase, but through a different mechanism. Instead of dephosphorylating AGE-1 itself, DAF-18 dephosphorylates the product of AGE-1’s catalysis, PIP_3_ (Ogg and Ruvkun 1998). DAF-18 is uniquely positioned to oppose insulin signaling by intersecting with one of the first steps of the pathway. Since neuroendocrine signals, including insulins, limit lifespan, we hypothesized that counteracting these signals may be crucial for the age-dependent activation of DAF-16, and so we investigated the role of DAF-18 in regulating DAF-16 during aging in *C. elegans*.

DAF-18 is the *C. elegans* ortholog of mammalian PTEN, a tumor suppressor that is frequently mutated in sporadic cancers (Milella *et al*. 2015). Although it functions primarily as a lipid phosphatase, PTEN can also dephosphorylate protein substrates. When localized to the cytosol, PTEN acts to inhibit processes that are of particular relevance to cancer, including progression through the cell cycle, cell growth, and cell migration. PTEN can also localize to the nucleus where it contributes to genome integrity by activating DNA repair pathways, and it plays a role in chromatin remodeling (Milella *et al*. 2015). Similar to PTEN, DAF-18 also inhibits cell proliferation primarily by depleting PIP_3_ levels and thus negatively regulating AKT-1. For example, during L1 diapause DAF-18 prevents germ cell proliferation that is mediated by AKT activation of the TOR complex and it also blocks division of Q neuroblasts promoted by AKT activation of MPK-1 (Fukuyama *et al*. 2006, 2012; Watanabe *et al*. 2008; Zheng *et al*. 2018). In the germline, DAF-18 activates the serine/threonine kinase PAR-4 to inhibit oocyte growth, maturation, and ovulation (Narbonne *et al*. 2015, 2017). In this capacity, DAF-18 acts locally within one gonad arm to oppose systemic IIS signaling that promotes oocyte growth. Notably, in none of the aforementioned roles does DAF-18 act by regulating DAF-16.

While capable of acting independently of DAF-16, prior to our study several lines of evidence suggested that DAF-18 plays a key role in determining lifespan by functioning through DAF-16. First, mutations in *daf-18* suppress the extended lifespan of *daf-2* mutants, similar to mutations in *daf-16* (Dorman *et al*. 1995). Along those lines, restoring *daf-18* expression in *daf-2(e1370);daf-18(mg198)* mutants rescues the lifespan extension phenotype of *daf-2* mutants in a *daf-16*-dependent manner (Masse *et al*. 2005). Third, *daf-18* is necessary for the enrichment in nuclear localization of DAF-16 in *daf-2* mutants (Larsen *et al*. 1995; Dorman *et al*. 1995; Lin *et al*. 2001). Finally, both *daf-16* and *daf-18* are required for lifespan extension through hormetic heat conditioning (Cypser *et al*. 2006). Taken together, these results strongly suggest that DAF-18 plays a role in aging, but specific functions for DAF-18 in adult animals have been established only in axon regeneration and associative learning (Tomioka *et al*. 2006; Byrne *et al*. 2014).

Here we show that the age-dependent increase in DAF-16 transcriptional activity requires DAF-18. We find substantial functional overlap between DAF-18 and DAF-16 during aging, suggesting that the two proteins act in concert. In particular, in postreproductive *C. elegans* both proteins contribute to innate immunity. Each confers resistance to microbial infection by functioning during adulthood in both the neurons and the intestine. DAF-18 appears to be expressed in chemosensory amphid neurons in adult *C. elegans*. Combined with our functional data, this suggests that it may be part of a neuroendocrine signaling axis between the neurons and the intestine to coordinate DAF-16 function as animals age. Finally, we uncover evidence to suggest that during aging DAF-18 may be deubiquitinated and sumoylated, two modifications that would promote its association with the plasma membrane to anchor it near its lipid substrates.

## MATERIALS AND METHODS

### *C. elegans* strains and maintenance

Worms were maintained using standard techniques as previously described (Brenner 1974). The *C. elegans* strains used in this study are as follows: Bristol wildtype N2, GR1329 [*daf-16(mgDf47)*], RB712 [*daf-18(ok480)*], VIL001 *mjyIs001 [plys-7::GFP]*, DAF-18::GFP zhEx343 [*daf-18 genomic::gfp*]; pCFJ90 [P*myo-2*::mCherry], NR350 [*rde-1(ne219); kzls20* [*hlh1*p::*rde-1* + *sur-5*p::NLS::GFP]], VP303 [*rde-1(ne219)*; *kbIs7* [*nhx-2*p::*rde-1* + *rol-6(su1006)*]], TU3401 [*sid-1(pk3321)*; *uIs69* [pCFJ90 (*myo-2*p::mCherry) + *unc-119*p::*sid-1*]]. To avoid possible confounding effects from UV-induced lesions at loci other than *daf-18*, RB712 was outcrossed 3 times prior to functional characterization

### Generation of age-matched cohorts of *C. elegans*

Synchronized populations of worms were obtained by sodium hypochlorite treatment of gravid adult animals to harvest eggs followed by hatching L1 larvae in M9 buffer in the absence of a bacterial food source for 16-20h at 22°C. Approximately 2000 L1 larvae were plated on to NGM media seeded with either *E. coli* OP50 *or E. coli* HT115 strains harboring RNAi constructs, depending on the experiment. To age animals to Day 6 of adulthood, worms were allowed to develop to the L4 larval stage before transferring them to new NGM plates containing 25 ug/mL 5-fluorodeoxyuridine (FUdR) and seeded with *E. coli*.

### RNAi treatment

*E. coli* HT115 strains carrying plasmids encoding dsRNA were thawed from glycerol stocks stored at −80°C and grown overnight at 37°C on LB plates supplemented with ampicillin and tetracycline. Single colonies were used to inoculate 200 mL of LB + ampicillin and incubated with shaking overnight at 37°C. To concentrate cells, cultures were centrifuged for 10 minutes at 3000x*g* and the cell pellet was resuspended in 20 mL of fresh ampicillin-supplemented LB. Approximately 1 mL of the concentrated cell cultures was used to seed two sets of NGM plates: one containing carbenicillin, and 2M IPTG (referred to as “RNAi plates”) and a second set containing carbenicillin, 2M IPTG, and 25ug/mL FUdR; referred to as “RNAi + FUdR plates”). Seeded plates were incubated in the dark at 22°C for at least 3 days before animals were added to them for knockdown experiments.

Synchronized populations of worms were subjected to RNAi treatments at one of two points during aging, depending on the experiment. For experiments involving a brief pulse of RNAi to knockdown *daf-18*, *daf-16*, or *age-1*, wild type N2 L1 larvae harvested from hypochlorite treatment of gravid adult hermaphrodites were maintained on NGM plates with lawns of *E. coli* OP50 at 20°C as described above until Day 4 of adulthood. Approximately 48 hours prior to challenging these animals with *P. aeruginosa*, they were transferred to RNAi +FUdR plates seeded with the *daf-18*, *daf-16*, or *age-1* RNAi clone (or L4440 vector control clones) and incubated at 20°C. For all other RNAi experiments, after hatching overnight in M9 buffer wild type N2 L1 larvae harvested from hypochlorite treatment of gravid adult hermaphrodites were cultivated on RNAi plates seeded with concentrated *E. coli* RNAi and maintained at 20°C. Upon reaching the L4 stage, these animals were transferred by chunking to RNAi + FUdR plates where they were separated from the agar chunk before it was removed from the plate. *C. elegans* were aged on these plates at 20°C until Day 6 of adulthood when they were harvested or used for other assays.

Transgenic strains of *C. elegans* were used to knock down *daf-18* and *daf-16* in a tissue-specific manner. In particular, to target these genes by RNAi in only the intestine or the muscle, in an *rde-1(ne219)* mutant background *rde-1* is supplied in an integrated transgene that drives its expression by a promoter that is specific to either the intestine (strain VP303) or the muscle (strain NR350). For neuron-specific RNAi, *sid-1* expression is restored only in neurons under the *unc-119* promoter in a *sid-1(pk3321)* background (strain TU3401).

### Fluorescence microscopy

To monitor the expression of *Plys-7::GFP* during aging, worms on RNAi or RNAi + FUdR plates were inspected under a Zeiss Stemi SV11 APO Stereo Microscope equipped with a Zeiss digital camera. In each of three independent biological replicates at least 200 animals spread over 4 separate plates per condition were scored for *Plys-7::GFP* expression levels at each time point. Depending on their pattern of fluorescence animals were assigned to one of three categories, as follows: 1) High—bright GFP signal observed throughout entire length of intestine, 2) Medium—comparatively weaker GFP signal that tended to be brighter in the anterior intestine than in the posterior intestine, 3) Low—dim GFP signal in all or only a segment of the intestine. To generate micrographs of the expression of the *Plys-7::GFP* reporter during aging, *C. elegans* maintained on NGM plates seeded with OP50 or subjected to feeding-based RNAi were mounted onto agarose pads (10% agarose in M9 buffer) seated on glass slides and immobilized in a slurry of 0.1 ìm polystyrene beads (Polysciences, Warrington, PA).

The expression of DAF-18::GFP was assessed by confocal microscopy using a Leica TCS SP8 inverted confocal microscope equipped with both HyD and standard PMTs (Leica). Day 6 adult *C. elegans* that had been maintained on NGM and (after reaching the L4 stage) NGM + FUdR plates seeded with *E. coli* OP50 were mounted on to agarose pads on glass slides as described above. Both the 488 nm and 552 nm laser lines were used to illuminate samples during image acquisition. Z-stacks consisting of 25-40 optical sections were collected for each region of interest at a resolution of 1024×1024 pixels at a magnification of either 400 or 630x. Composite images were initially processed with Leica LASX software, and further adjustments were made using NIH ImageJ.

### Pseudomonas aeruginosa infection

Larval stage and adult *C. elegans* were infected with the bacterial pathogen *Pseudomonas aeruginosa* as described (Youngman *et al*. 2011). Briefly, approximately 100 animals of the indicated ages were transferred from plates containing *E. coli* OP50 or *E. coli* HT115 (for RNAi treatments) and distributed as three groups of ∼30 worms each onto plates seeded with *P. aeruginosa* strain PA14 which were then incubated at 25°C. At intervals of 12-24 hours, the survival of *C. elegans* on the *P. aeruginosa* plates was assessed and dead animals were removed at each time point until all of the worms in the assay had died. At least three independent biological replicates of each infection experiment were performed.

### Calculation of median survival times and statistical analysis

To calculate the median survival time (LT_50_) of *C. elegans* infected with *P. aeruginosa*, the fraction of animals alive at each time point during an assay was first plotted as a function of time in Excel, accounting for animals that were inadvertently overlooked or that escaped between time points. These data were then imported into SigmaPlot (Systat Software, San Jose, CA) and a three parameter sigmoidal curve was fit according to the general equation y=a/(1+e^(-(x-x0)/b)^). This equation was used to determine the point at which 50% of the animals in the assay had died (Efron 1987). The average fold difference between the LT_50_ of mutant strains or experimental RNAi treatments and control animals was calculated, and the statistical significance of that difference was assessed using a one-sample t-test after first applying a Shaprio-Wilk test for normality.

### RNA isolation

To harvest *C. elegans* for RNAi isolation, approximately 10,000-20,000 animals were washed off of NGM plates in M9 buffer and allowed to settle to the bottom of a 15 mL conical tube by gravity. After removing the supernatant the worms were washed with 10 mL of fresh M9 and again allowed to settle to the bottom of the tube. All but 1-2 mL of buffer were removed and used to resuspend the worms before transferring them to cryovials where more buffer was removed, leaving behind sufficient volume to cover the worm pellet. After adding 300 ìL of Trizol reagent (ThermoFisher Scientific, Waltham, MA), the worms were vortexed in a series of 30 second intervals over a period of 5 minutes interspersed by brief rest periods and then transferred to −80°C for storage. To prepare total RNA from worm pellets, frozen animals in Trizol were thawed at room temperature and then briefly vortexed before pelleting in a microcentrifuge at 16,000x*g* for 5 min to remove worm carcasses and debris. Following a phenol/chloroform extraction, total RNA was precipitated in isopropanol, washed in 70% ethanol, and resuspended in RNase-free H_2_O. RNA was then reisolated over a column using the RNeasy kit (Qiagen, Germantown, MD). The absorbance of samples at 260 nm was measured by Nanodrop (ThermoFisher Scientific, Waltham, MA) and used to calculate the concentration of RNA.

### qRT-PCR

To measure gene expression levels by qRT-PCR, cDNA was reverse transcribed from 1 ìg samples of total RNA using Superscript IV Vilo RT(Invitrogen). Diluted cDNA was used as a template in PCR reactions carried out in Power SYBR Green master mix (Applied Biosystems, Foster City, CA). Previously described primer pairs (Troemel *et al*. 2006) were used to amplify sequences from *lys-7*, *sod-3*, *mtl-1,* and *cpr-2* transcripts with *tba-1* as the reference, and fluorescence was quantified using BioRad CFX96 thermocycler (BioRad, Hercules, CA). Relative gene expression levels were calculated using the ΔΔCt method (Livak and Schmittgen 2001). This analysis was performed on a total of three biological replicates, representing three independently generated cohorts of aging worms.

### Data availability

Files uploaded as supplemental material to figshare are described below. Table S1 contains the raw data from manual scoring of *plys-7::GFP* expression in three independent biological replicates corresponding to the data shown as bar graphs in Fig. 2 B-D. A separate section of the table shows the raw scores converted to fractions so that the proportion of animals in each category may be compared. Table S2 lists raw Ct values and calculated ΔΔCt for *mtl-1* and *sod-3* expression levels in L4 larvae and Day 6 adults treated with RNAi targeting *daf-18*, *daf-16*, or *age-1*. These data correspond to the bar graph in Fig. 2E. Table S3 lists raw Ct values and calculated ΔΔCt for *cpr-2* and *lys-7* expression levels in L4 larvae and Day 6 adults treated with RNAi targeting *daf-18*, *daf-16*, or *age-1*. These data correspond to the bar graph in Fig. 2E. Table S4 lists compiled fold differences in expression of *mtl-1*, *sod-3*, *lys-7*, and *cpr-2* between L4 and Day 6 in animals treated with RNAi targeting *daf-16, daf-18*, or *age-1* for three independent biological replicates. Also listed are average fold differences in expression between Day 6 and L4 with standard deviations and standard error. These data correspond to the bar graph in Fig. 2E. Table 5 lists experimental details and statistical analyses corresponding to each survival curve in Figs. 3-5. Figure S1 shows confocal micrographs of *daf-18::GFP*-expressing worms treated with RNAi targeting GFP. Figure S2 shows additional replicates of the *P. aeruginosa* infection assay. Figure S3 shows additional replicates of the RNAi experiment to investigate the timing requirement of *daf-18* in *C. elegans* immunity. Figures S4 and S5 depict results from two additional replicates of the experiment in which RNAi was used to knock down *daf-18* expression in specific tissues before worms were infected with *P. aeruginosa.* Figure S6 shows an additional replicate of the experiment in which worms treated with RNAi targeting candidate DAF-18 regulators were infected with *P. aeruginosa*.

## RESULTS

### Expression pattern of DAF-18 in adult *C. elegans*

In *C. elegans* larvae a *daf-18* transcriptional reporter is expressed in a number of tissues, including the intestine, body wall muscles, seam cells, hypodermis, and neurons (Masse *et al*. 2005). The expression of endogenous DAF-18 in wildtype animals is most evident in the Z2/Z3 germline precursor cells and in oocytes, yet when overexpressed or in a *daf-2* mutant background, DAF-18 is detected in the intestine, the ventral nerve cord, and in head amphid neurons (Brisbin *et al*. 2009; Liu *et al*. 2014). We asked whether DAF-18 is still found in these same tissues in postreproductive adult *C. elegans* and so we examined the expression of DAF-18::GFP in adult animals six days after they had reached the L4 larval stage (hereafter referred to as Day 6 adults). At lower magnification, DAF-18::GFP signal was observed in two locations: near the procorpus and terminal bulb of the pharynx and, more faintly, within two broad parallel segments near the animal’s posterior (Fig. 1A). The substantial amount of autofluorescence in the intestine from lipofuscin precluded our ability to make conclusive observations regarding the expression of DAF-18 in that tissue in adult *C. elegans*. Closer examination at higher magnification revealed that DAF-18::GFP appears to be expressed in head neurons whose dendrites traverse the isthmus of the pharynx and extend toward the mouth, reminiscent of its expression in amphid neurons in younger animals (Fig. 1B; (Brisbin *et al*. 2009)). Surprisingly, at the posterior, what appeared as a diffuse pattern at lower magnification was actually discrete rows of puncta. While the specific identity of these structures awaits further confirmation, we note that their pattern is strikingly similar to the appearance of fragmented mitochondria in the body wall muscle of animals subjected to acute thermal stress (Fig. 1C (Momma *et al*. 2017)). Our imaging studies show that in postreproductive adult *C. elegans* DAF-18 continues to be expressed in many of the same tissues in which it is found in younger animals, suggesting that its role in those tissues is not limited to developmental processes or other functions that are exclusive to larvae.

**Figure 1.**
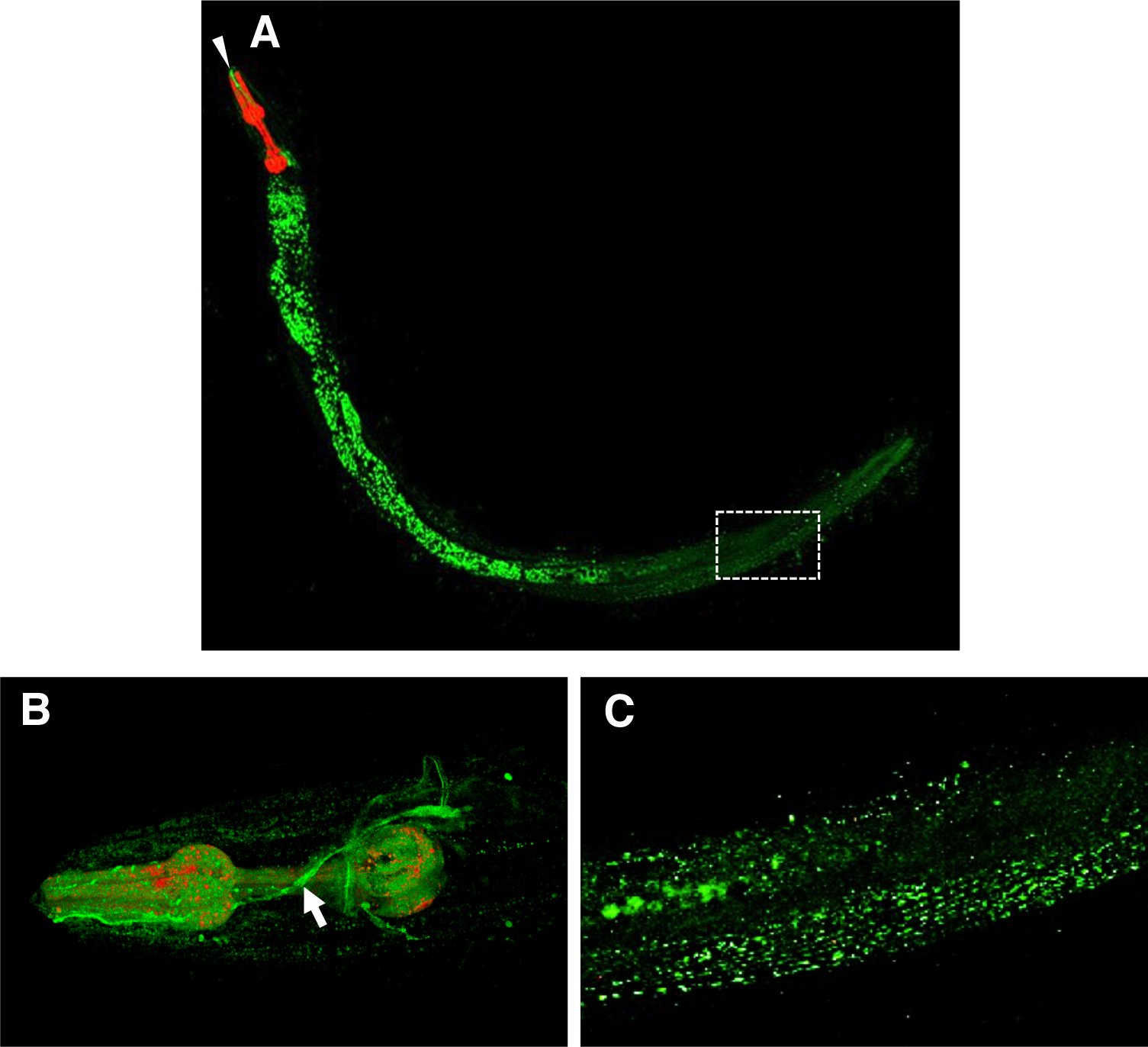
**DAF-18 is primarily expressed in head neurons in adult *C. elegans.*** Day 6 adult animals of strain zhEx343 expressing DAF-18::GFP were imaged by confocal microscopy. (A) Autofluorescence from lipofuscin in the intestinal cells is the predominant signal in images of whole adult animals. Comparatively weaker GFP expression is observed in the head (arrowhead) near the pharynx (red) and toward the posterior. (B) Higher magnification of the head region reveals that DAF-18::GFP is expressed in a presumptive amphid neuron whose dendrite crosses the isthmus of the pharynx (arrow) and extends to the mouth. (C) Higher magnification of the region bounded by the rectangle in (A) shows DAF-18::GFP expression in body wall muscle.

**Figure 2.**
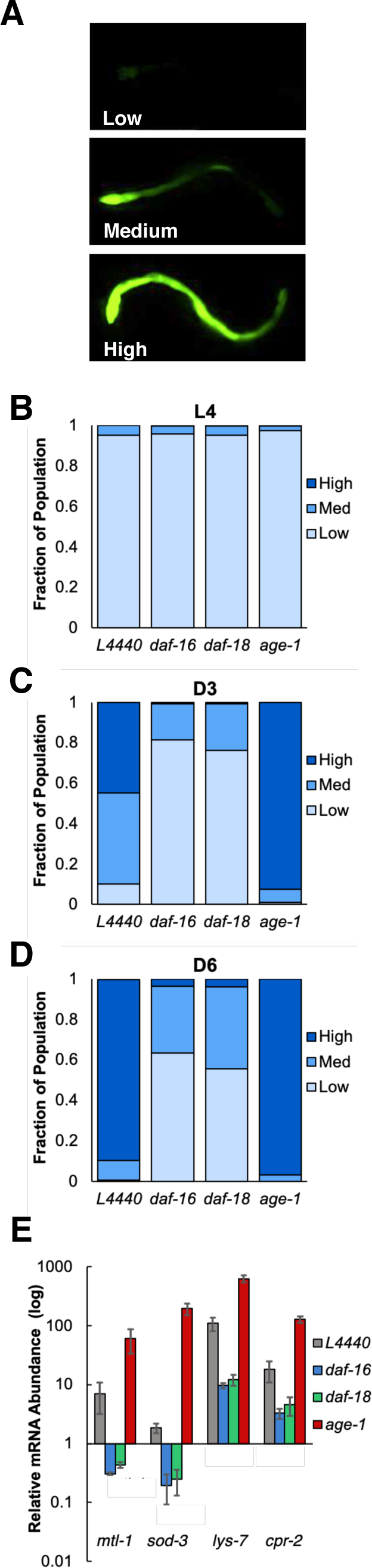
**The age-dependent increase in the transcriptional activity of DAF-16 requires DAF-18. (**A) Expression of the *Plys-7::GFP* reporter in the intestine of Day 6 adult *C. elegans*. Based on the expression level of Plys-7::GFP, individual animals within a population may be classified as either low, medium or high expressers. Micrographs show representative animals belonging to each category. (B-D) N2 wild type animals treated with an empty RNAi vector (L4440) or with RNAi targeting *daf-16*, *daf-18*, or *age-1* were scored according to the degree of *Plys-7::GFP* expression at the L4 stage (B), Day 3 of adulthood (C) or Day 6 of adulthood (D). At each time point, bars corresponding to a particular RNAi treatment are color coded to represent the fraction of the population scored as having high (green), medium (blue), or low (black) *Plys-7::GFP* expression. Shown is the average number of animals in each category over three independent biological replicates. In each replicate at least 200 animals from each RNAi treatment group were scored. (E) Endogenous transcript levels of DAF-16 transcriptional targets *mtl-1*, *sod-3*, *lys-7*, and *cpr-2* were measured in Day 6 N2 wild type animals treated with an empty RNAi vector (L4440) or with RNAi targeting *daf-16*, *daf-18*, or *age-1*. The mRNA abundance of each transcript was normalized to the levels of the housekeeping gene *tba-1* and is reported relative to its abundance in L4 larvae subjected to the same RNAI treatments. Error bars, standard deviation.

### DAF-18 is required for the age-dependent increase in the transcriptional activity of DAF-16

Similar to the effect of mutations in *daf-16*, both reduced-function and loss-of-function mutations in *daf-18* suppress the constitutive dauer and extended lifespan phenotypes of *daf-2* mutants (Dorman *et al*. 1995). This suggests that even in the absence of a repressive signal through the insulin receptor, DAF-18 is necessary for the full activation of DAF-16, which is responsible for the stress resistance and exceptional longevity of *daf-2* animals (Ogg and Ruvkun 1998). In wildtype larvae, DAF-16 appears to only be activated in response to acute environmental insults, remaining sequestered in the cytosol under nutritionally replete, stress-free conditions (McHugh *et al*. 2020). Shortly after animals transition to adulthood, however, *daf-16* expression is upregulated despite the absence of overt stressors (Bansal *et al*. 2014). This is met with a commensurate increase in DAF-16 transcriptional activity toward a subset of its targets, including those that confer resistance to bacterial pathogens (Li *et al*. 2019; McHugh *et al*. 2020). To determine whether DAF-18 regulates the age-dependent increase in DAF-16 transcriptional activity, we first asked whether DAF-18 was necessary for the increased expression of *plys-7::GFP*, an *in vivo* reporter in which the promoter of the DAF-16 transcriptional target gene *lys-7* drives GFP expression in the intestine (Alper *et al*. 2007).

The expression of *plys-7::GFP* increases in an age-dependent manner, and this requires *daf-16* (McHugh *et al*. 2020). As previously reported, we observed three major categories of *plys-7::GFP* expression levels within the intestines of age-matched cohorts of *C. elegans* (Fig. 2A). Animals in the “low” category showed weak *plys-7::GFP* expression that appeared to be confined to only the anterior-most portion of the intestine. “Medium” levels of *plys-7::GFP* expression were characterized by prominent GFP signal in the anterior intestine with weaker expression in posterior segments. *C. elegans* with robust *plys-7::GFP* expression throughout the entire length of the intestine without interruption were classified as “high” expressors. To test the possible role for DAF-18 in activating DAF-16 during aging, we examined the effect of knocking down *daf-18* on the expression of *plys-7::GFP* expression at three points: the L4 larval stage, Day 3 of adulthood, and Day 6 of adulthood. For comparison, *daf-16* and *age-1* were also targeted by RNAi. At each time point, animals were examined by fluorescence microscopy and were scored as belonging to one of the three categories described above, according to the level of *plys-7::GFP* expression in their intestine. If DAF-16 is regulated by DAF-18 during aging, we expected that RNAi targeting *daf-18* would have the same effect on *plys-7::GFP* expression during adulthood as knocking down *daf-16*. This is, in fact, what we found. In L4 larvae, nearly all animals (more than 90% in all cases) exhibited low expression *of plys-7::GFP*, regardless of what gene was targeted by RNAi (Fig. 2B). By Day 3 of adulthood, *plys-7::GFP* expression had increased in control animals fed *E. coli* harboring an empty RNAi vector (L4440), with animals being split almost evenly between the “high” and “low” expression categories (Fig. 2C). There was not an equivalent increase in *plys-7::GFP* expression when either *daf-18* or *daf-16* was knocked down. Instead, while medium levels of *plys-7::GFP* expression were observed in about 30% of animals, the majority of worms in both of these treatment groups still showed low levels of *plys-7::GFP* expression. By contrast, RNAi targeting *age-1* enhanced the age-dependent increase in *plys-7::GFP* expression, resulting in a larger proportion of Day 3 *C. elegans* being assigned to the “high” category. This same general pattern persisted through Day 6 of adulthood. At that point, almost all *C. elegans* in the control group showed high levels of *plys-7::GFP* expression, similar to animals treated with RNAi targeting *age-1* (Fig. 2D). RNAi of *daf-18* and *daf-16* largely suppressed the increased expression of *plys-7::GFP* in Day 6 animals. Compared to L4 larvae, the fraction of Day 6 animals with medium levels of *plys-7::GFP* expression still increased when *daf-18* or *daf-16* were knocked down, yet fewer than 10% had high *plys-7::GFP* expression levels and in about half of the animals its expression was low. Together, the results of our *in vivo* reporter assay suggest that DAF-18 is necessary for the activation of DAF-16 transcriptional activity that occurs during aging in *C. elegans*.

To validate the results of our studies with *plys-7::GFP*, we measured the levels of endogenous transcripts of *lys-7* along with the transcripts of three other genes whose expression is regulated by DAF-16, *mtl-1*, *sod-3*, and *cpr-2*. (Troemel *et al*. 2006; Kwon *et al*. 2010). qRT-PCR was used to compare the expression levels of the four DAF-16 transcriptional targets in L4 larvae to their expression levels in Day 6 adults (Fig. 2E). Each of the genes that we tested was more highly expressed in adult *C. elegans* than in larvae, although the magnitude by which they were upregulated differed. For example, *lys-7* was upregulated by almost 100-fold in Day 6 adults compared to L4 animals, while the expression of *sod-3* was only about 2-fold greater. To test whether *daf-18* is necessary to bring about the changes in expression of DAF-16 transcriptional targets over time that we observed, we used RNAi to knock down its expression and evaluated the effect relative to knocking down *daf-16* or *age-1*. Knockdown of *daf-18* caused substantial reductions in the expression of DAF-16 targets, phenocopying the effect of knocking down *daf-16* (Fig. 2E). Targeting either *daf-18* or *daf-16* with RNAi decreased the expression of *mtl-1* and *sod-3* in Day 6 adults to levels that were lower than at the L4 larval stage and reduced *lys-7* and *cpr-2* expression by nearly 10-fold. In contrast, knocking down *age-1* increased the expression of the DAF-16 transcriptional targets in Day 6 *C. elegans* by two to three orders of magnitude. These results are consistent with our observations with the *plys-7::GFP* reporter, providing additional evidence that DAF-18 regulates the age-dependent increase in DAF-16 transcriptional activity. Further, they imply that DAF-18 and AGE-1 play opposing roles during aging in wildtype *C. elegans*.

### DAF-18 functions in innate immunity in adult *C. elegans*

We previously reported that one functional consequence of the age-dependent increase in DAF-16 transcriptional activity is that DAF-16 contributes to innate immunity in adult but not larval stage *C. elegans* (McHugh *et al*. 2020). Since we found that in adults the elevated levels of two DAF-16 targets with potential functions as immune effectors, *lys-7* and *cpr-2*, require DAF-18 we asked whether DAF-18 contributes to host defense during aging. To test this possibility, the survival of *daf-18(ok480)* mutants infected with the human opportunistic pathogen *Pseudomonas aeruginosa* at either the L4 larval stage or at Day 6 of adulthood was compared to the survival of age-matched infected *daf-16(mgDf47)* or N2 wild type animals. When challenged with *P. aeruginosa* as L4 larvae, *daf-18(ok480)* mutants died at a similar rate as wildtype animals, as did *daf-16(mgDf47)* mutants (Fig. 3A; Table S5). On the other hand, when the infection was initiated at Day 6 of adulthood, *daf-18(ok480)* mutants and *daf-16(mgDf47)* animals both died more rapidly than wildtype worms (Fig. 3B; Table S5). Stated in more quantitative terms, the infected adult *daf-18(ok480)* and *daf-16(mgDf47)* mutants had median lifespans (LT_50_) of ∼70 hours, whereas the LT_50_ of infected N2 wildtype adults was ∼120 hours. These data indicate that, similar to DAF-16, DAF-18 functions in innate immunity in adult *C. elegans*.

**Figure 3.**
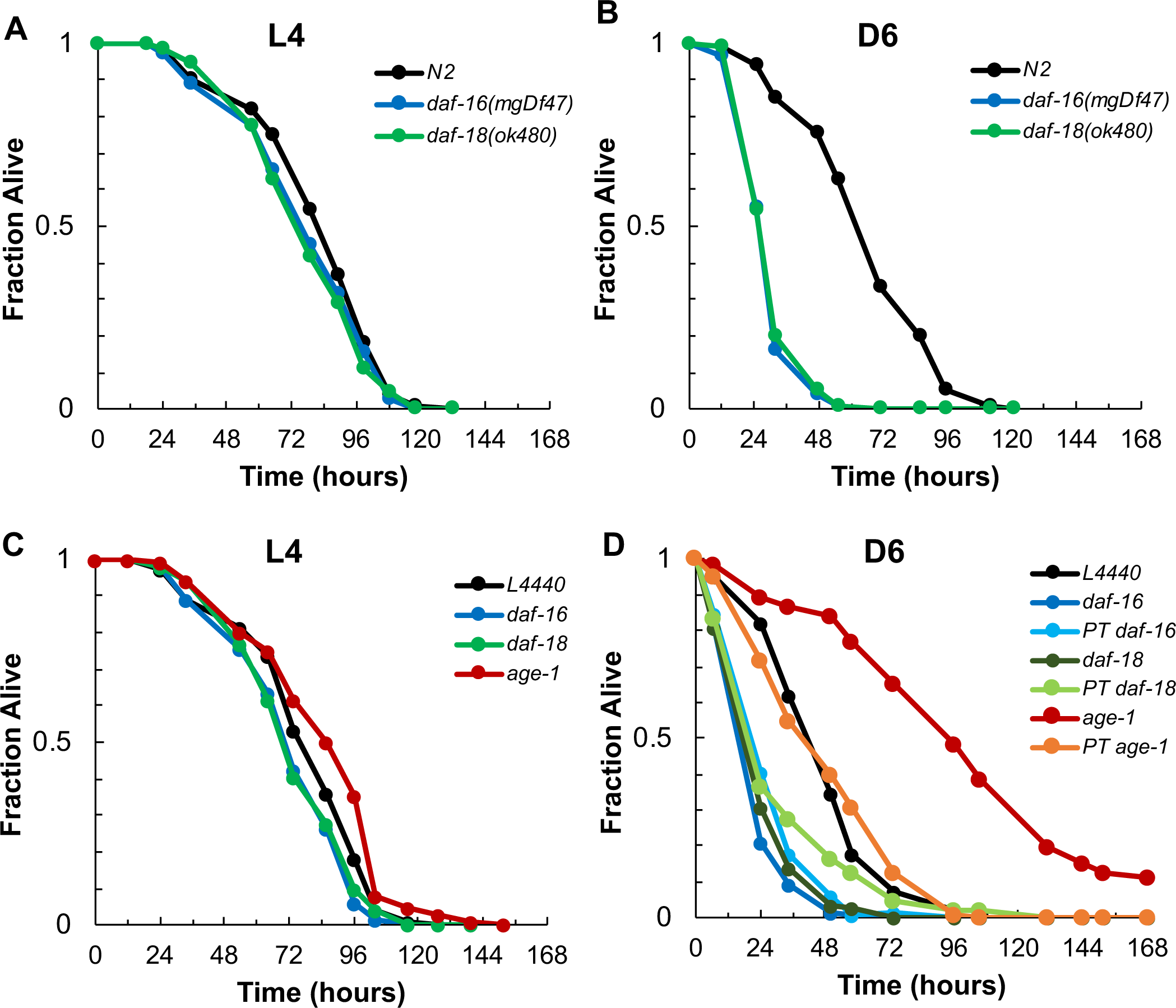
DAF-18 contributes to innate immunity during adulthood in *C. elegans*. (A and B) Age-matched cohorts of N2 wildtype animals, *daf-16(mgDf47)*, or *daf-18(ok480)* mutants were infected with *P. aeruginosa* at the L4 larval stage (A) or at Day 6 of adulthood (B) and their survival was recorded over time. Plotted in each panel is the fraction of animals alive as a function of time after the infection was initiated. (C and D) Beginning at the L1 larval stage N2 wildtype animals were treated with an empty RNAi vector (L4440, black) or with RNAi targeting *daf-16* (dark blue), *daf-18* (dark green), or *age-1* (red) and then infected with *P. aeruginosa* at either the L4 larval stage (C) or at Day 6 of adulthood (D). In another group of animals, RNAi knocking down *daf-16* (light blue), *daf-18* (light green), or *age-1* (orange) was initiated at Day 4 and worms were infected with *P. aeruginosa* at Day 6 (D). Plotted in each panel is the fraction of animals alive as a function of time after the start of the infection.

That *daf-18(ok480)* phenocopies the enhanced pathogen susceptibility of *daf-16(mgDf47)* implies that DAF-18 cooperates with DAF-16 to confer resistance to infection during aging. If this is the case, then both genes should be functioning in innate immunity simultaneously. Said another way, the timing requirements for *daf-18* and *daf-16* in host defense should be the same. While it is ubiquitously expressed during larval development and for the duration of life in *C. elegans*, DAF-16 functions exclusively during adulthood to fulfill its roles in innate immunity and in determining lifespan (Dillin *et al*. 2002; McHugh *et al*. 2020). For these two aspects of DAF-16’s function its temporal requirement was defined by initiating RNAi to reduce its expression only during discrete intervals instead of according to more commonly used regimens that involve continuous knockdown, beginning at an early larval stage or in the parental generation.

We employed a similar strategy to determine the timing requirement of DAF-18 in host defense. Specifically, we compared the effect of sustained RNAi treatment that knocked down *daf-18* expression starting at the L1 larval stage and continuing through adulthood to a brief treatment in which RNAi was delayed until Day 4 of adulthood. Animals treated according to the latter scheme would therefore be allowed to develop and complete the reproductive period in the presence of normal *daf-18* expression levels. If the timing requirements for DAF-18 and DAF-16 were the same, then we anticipated that waiting until Day 4 of adulthood to knock down *daf-18* expression would be sufficient to recapitulate the enhanced pathogen susceptibility phenotype that we observed when *daf-18* RNAi treatment began at L1, just as it is for bringing about the *daf-16* pathogen susceptibility phenotype (McHugh *et al*. 2020).

We found that the sustained RNAi treatment accurately reproduced the phenotype of the *daf-18(ok480)* loss-of-function mutant. That is, knockdown of *daf-18* from the L1 stage did not cause L4 larvae to be more susceptible to *P. aeruginosa* infection than untreated control (L4440) animals (Fig. 3C). At Day 6 of adulthood, however, animals subjected to this RNAi regimen targeting *daf-18* died from bacterial infection at rate that was similar to when the RNAi was directed against *daf-16*, and both sets of *C. elegans* died faster than infected control worms treated with an empty RNAi vector (Fig. 3D). Delivering just a short pulse of RNAi only to adult worms turned out to have this same effect. When *daf-18* was knocked down for two days during adulthood, from Day 4 until *P. aeruginosa* infection was initiated at Day 6, animals were still more susceptible to the infection than untreated controls and died at a similar rate to worms subjected to the sustained RNAi treatment (Fig 3D). Moreover, the LT_50_ of Day 6 animals subjected to the brief RNAi treatment targeting *daf-18* closely approximated the LT_50_ of animals in which *daf-16* was knocked down using the same regimen, corroborating our previous observations (Fig. 3D; (McHugh *et al*. 2020)). These results indicate that DAF-18 and DAF-16 have identical timing requirements for their roles in innate immunity and that their contributions to host defense are not the products of having acted earlier in the worm’s life but instead are made at the point just prior to when animals were challenged with *P. aeruginosa*. That is, both proteins function during adulthood to protect adult *C. elegans* from bacterial infection.

Within the IIS pathway, DAF-18 counteracts the function of the AGE-1 kinase that is immediately downstream of the DAF-2 insulin receptor by dephosphorylating PIP_3_, thus depriving PDK-1 of a lipid that promotes its activation. Considering the antagonistic relationship between DAF-18 and AGE-1, abrogating the function of one protein should have the opposite effect of when the function of the other is compromised. This is what we observed in our analysis of DAF-16 transcriptional target expression levels in Day 6 adults when *daf-18* or *age-1* were knocked down (Fig. 2D, E). Based on those results, we predicted that whereas *daf-18* promotes innate immunity during aging, *age-1* might act to inhibit it. To test this possibility, we subjected worms to sustained RNAi treatment targeting *age-1* beginning at the L1 stage of larval development. When challenged with *P. aeruginosa* as L4 larvae, the survival of these animals was similar to that of untreated control worms (Fig. 3C). Yet when the infection was initiated at Day 6 of adulthood, *age-1* knockdown caused worms to become more resistant to the pathogen, extending the median survival of infected *C. elegans* by nearly two-fold (Fig. 3D; Table S5). This result confirmed our hypothesis and suggests that *age-1* functions to limit host defense in postreproductive animals but not in larvae. When we investigated the timing requirement for *age-1* by treating worms with RNAi to knock down *age-1* beginning at Day 4 of adulthood before infecting them with *P. aeruginosa* two days later, these animals died at the same rate as untreated controls, suggesting that AGE-1 functions prior to Day 4 to regulate host defense in adults (Fig. 3D).

### DAF-18 and DAF-16 function in the intestine and neurons to protect adult *C. elegans* from bacterial pathogens

Our functional characterization of DAF-18 during aging supports the possibility that it acts together with DAF-16 to modulate innate immunity in adult *C. elegans*. Since DAF-16 is downstream of DAF-18 in the IIS pathway, this could mean that the two proteins function in the same tissues. Although it is expressed in many other tissues, the sites of action of DAF-16 for its role in lifespan determination are the intestine and neurons (Wolkow *et al*. 2000; Libina *et al*. 2003). We asked about the site of action of both DAF-18 and DAF-16 for their roles in host defense by knocking down their expression only in individual tissue types through the use of tissue-specific RNAi strains of *C. elegans*. We expected that inhibiting *daf-16* or *daf-18* expression in the particular tissue in which they are required for innate immunity should be sufficient to recapitulate the effect of systemic RNAi that reduces their expression in all tissue types (Fig. 4A and B; Table S5). Although DAF-18::GFP appears to be expressed in body wall muscle (Fig. 1C), muscle-specific RNAi of *daf-18* had no effect on the susceptibility of either L4 larvae or Day 6 adults to *P. aeruginosa* infection (Fig. 4C and D). Similarly, knockdown of *daf-16* in the muscle did not affect the survival of animals infected as either L4 larvae or Day 6 adults. Inhibiting the expression of *daf-18* and *daf-16* in either the intestine or the neurons, however, caused Day 6 adults to be just as sensitive to *P. aeruginosa* infection as when *daf-18* or *daf-16* were knocked down in all tissues (Fig. 4E-H). These results indicate that DAF-18 and DAF-16 act in both the intestine and neurons in postreproductive adult worms and that their functions in those tissue promote host defense.

**Figure 4.**
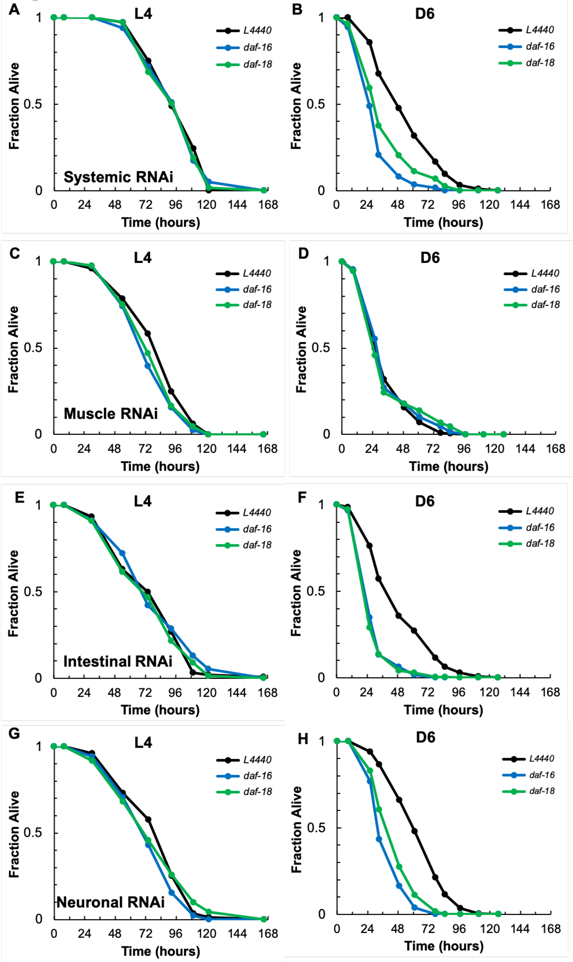
DAF-18 functions in the intestine and neurons of adult animals to protect them from bacterial pathogens. Beginning at the L1 larval stage N2 wild type animals (A and B) along with mutants in which RNAi-mediated gene knockdowns are restricted to the muscle (C and D), the intestine (E and F) or the neurons (G and H) were treated with an empty RNAi vector (L4440, black) or with constructs targeting *daf-18* (green) or *daf-16* (blue). Animals were infected with *P. aeruginosa* at either the L4 larval stage (A, C, E, G) or at Day 6 of adulthood (B, D, F, H) and their survival was recorded over time. In each panel, the fraction of worms alive is plotted as a function of time since the infection was initiated. The data shown are representative of three independent replicates.

### Posttranslational modifications modulate the activity of DAF-18 during aging

Similar to PTEN in mammals, DAF-18 is regulated at several levels in *C. elegans*. For example, the HLH-25 transcription factor inhibits *daf-18* expression and thus appears to be a functional ortholog of HES1, which negatively regulates PTEN in human cells and in mice (Chou *et al*. 2015). At the protein level, in general phosphorylation inhibits DAF-18 by decreasing its stability. For example, DAF-18 is phosphorylated by the Eph receptor tyrosine kinase VAB-1, and *vab-1* mutants have higher levels of DAF-18 protein than wildtype animals (Brisbin *et al*. 2009). Interestingly, a human PTEN allele with threonine-to-alanine substitutions that destroy phosphorylation sites was able to rescue the short lifespan phenotype of *daf-2(e1370);daf-18(mg198)* mutants more effectively than the wildtype PTEN allele, suggesting that the lack of phosphorylation made PTEN more active (Solari *et al*. 2005). The scenario during aging that is supported by our functional analysis implies that DAF-18 activation increases in adult *C. elegans*. Since most previous investigations of DAF-18 regulation have identified or characterized negative regulators, to search for potential DAF-18 activators, we selected *C. elegans* orthologs of three positive regulators of PTEN—*smo-1*, *math-33*, and *cpb-3*-- for functional analysis studies (Bermúdez Brito *et al*. 2015). *smo-1* encodes the sole *C. elegans* ortholog of mammalian SUMO, a ubiquitin-like moiety that regulates protein function. In mammals, sumoylation of PTEN influences it subcellular localization ((Huang *et al*. 2012). MATH-33 is an ortholog of human USP7 that deubiquitinates target proteins in a thiol-dependent manner. CBP-3 has predicted histone acetyltransferase activity and is an ortholog of human CREB binding protein (CBP). Depending on the specific nature of the modification, ubiquitination either destabilizes PTEN or promotes its localization to the nucleus and so MATH-33 would be expected to influence those same properties of DAF-18 (Chen *et al*. 2018). For comparison, we also characterized one putative negative regulator of DAF-18, SUP-17. With predicted metallopeptidase activity, SUP-17 is an ortholog of human ADAM10 which, by proteolytically activating Notch or receptor tyrosine kinases (such as HER2), may inhibit either the expression or catalytic activity of PTEN (Wen *et al*. 1997).

To determine whether our candidate genes might regulate DAF-18 during aging, we assessed their roles in host defense in L4 larvae and in Day 6 adults. We reasoned that knocking down genes that encode putative activators of DAF-18 would phenocopy the knockdown of *daf-18*, curtailing the survival of adult *C. elegans* infected with *P. aeruginosa*, but having no effect on the ability of L4 larvae to resist bacterial infection. Conversely, RNAi targeting potential DAF-18 inhibitors should lead to the opposite phenotype, conferring resistance to bacterial infection during adulthood. None of the genes that we tested appear to influence innate immunity in L4 worms, since RNAi treatments targeting each gene individually failed to modify the survival of animals infected at that stage of development (Fig. 5A, Table S5). When animals were challenged with *P. aeruginosa* at Day 6 of adulthood, RNAi targeting *cbp-3* did not produce a phenotype. Knockdown of *smo-1* or *math-33*, however, both reduced the median survival of infected animals to a similar extent as RNAi targeting *daf-18* or *daf-16* when compared to untreated animals (Fig. 5B). This agrees with our prediction for the phenotype of possible positive regulators of the age-dependent function of DAF-18, and it is consistent with the established roles for the orthologs of SMO-1 and MATH-33 in PTEN regulation. These data suggest that during aging in *C. elegans* the activity of DAF-18 may be regulated by post-translational modifications, including sumoylation and ubiquitination.

**Figure 5.**
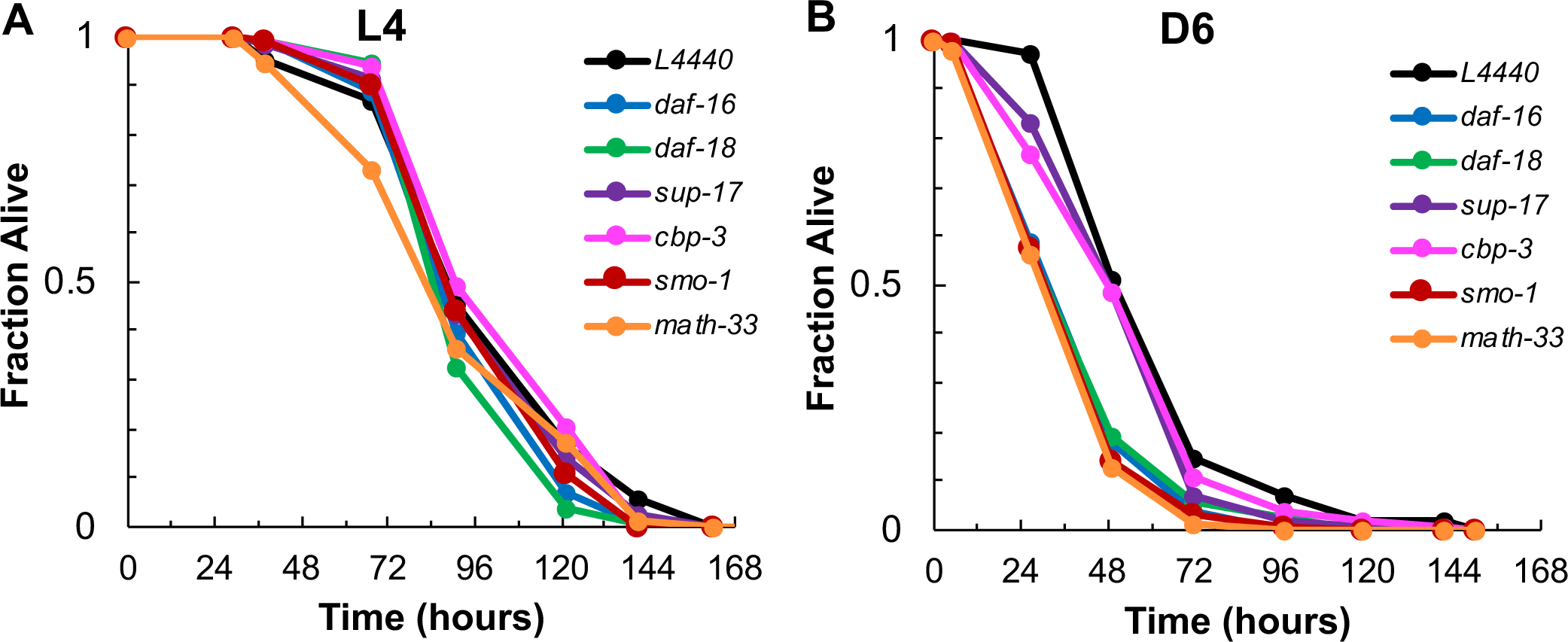
The age-dependent function of DAF-18 in innate immunity may be regulated by post-translational modifications. Following RNAi treatment beginning at the L1 larval stage to knock down the candidate DAF-18 regulators *sup-17* (purple), *cbp-3* (pink), *smo-1* (red), or *math-33* (orange), N2 wild type *C. elegans* were challenged with *P. aeruginosa* as either L4 larvae (A) or as Day 6 adults (B). Their survival was monitored over time and was compared to that of age-matched infected animals in which *daf-18* (green) or *daf-16* (blue) were targeted by RNAi or animals that were treated with an empty RNAi vector (L4440, black). In each panel, the fraction of worms alive is plotted as a function of time since the infection was initiated. The data shown are representative of three independent replicates.

## DISCUSSION

Considering its critical, evolutionarily conserved roles in lifespan determination and stress resistance, understanding the mechanism by which DAF-16 is regulated in *C.elegans* has the potential to reveal paradigms governing the activity of FOXO transcription factors in other species. While DAF-16 has been known to be activated by acute stress for some time, more recent evidence has revealed that the transcriptional activity of DAF-16 increases as animals age even without the trigger of an environmental insult (Li *et al*. 2019; McHugh *et al*. 2020). This implies that there is a regulatory mechanism inherent to the aging process that governs DAF-16 activity once animals reach adulthood. So far the only putative regulator of the age-dependent activation of DAF-16 to be identified is SMK-1, a component of the PP4 phosphoprotein phosphatase complex that, based on homology, modulates the activity of the catalytic subunit (McHugh *et al*. 2020). Here we tested the possibility that DAF-18, the *C. elegans* ortholog of mammalian PTEN that antagonizes PI3 kinase in the IIS pathway, also plays a role in regulating DAF-16 during adulthood. Although DAF-18 functions in adult animals to inhibit axon regeneration and associative learning, both processes are at least partially independent of DAF-16 (Tomioka *et al*. 2006; Byrne *et al*. 2014). Our data suggest an additional regulatory function for DAF-18 during aging that directly impacts the transcriptional activity of DAF-16 and, consequently, its role in innate immunity. We found that the age-dependent increase in expression of DAF-16 targets requires DAF-18 and that without *daf-18* host defense in postreproductive adult animals is compromised, enhancing their susceptibility to lethal infection by bacterial pathogens, just as the loss of functional *daf-16* does. Our results suggest that as *C. elegans* age an inhibitory signal through the IIS must be counteracted by DAF-18 in order to initiate a basal level of DAF-16 transcriptional activity that buffers older animals against anticipated but yet to be encountered environmental stress. We anticipate that our findings will have implications for PTEN-mediated regulation of aging processes in evolutionarily diverse species, including humans.

Prior to our work, a connection between *daf-18* and aging had been established by studies examining its role as a suppressor of the extended lifespan of *daf-2* mutants (Larsen *et al*. 1995; Dorman *et al*. 1995; Ogg and Ruvkun 1998; Mihaylova *et al*. 1999; Honda and Honda 2002; Patel *et al*. 2008). Significantly, in the absence of any other genetic lesions, mutations in *daf-18* shorten lifespan while *daf-18* overexpression extends lifespan, indicating that DAF-18 contributes to longevity when IIS signaling is intact in wildtype animals (Dorman *et al*. 1995; Mihaylova *et al*. 1999; Honda and Honda 2002; Masse *et al*. 2005; Brisbin *et al*. 2009). It is under these same conditions that DAF-16 becomes activated during the course of aging, and this is why we decided to investigate a possible role for DAF-18 in regulating DAF-16 during adulthood. A direct output of DAF-16 activation is a change in gene expression, and we measured this in two ways in adult animals in the presence and absence of DAF-18. Both our qualitative assessment of DAF-16 transcriptional target expression using the *in vivo plys-7::GFP* reporter and our quantitative measurement of endogenous transcript levels showed that increased DAF-16 transcriptional activity during adulthood depends on DAF-18. While transcriptomic analysis would provide a more complete picture of the full extent of DAF-18’s influence on the transcriptional output of DAF-16, we found DAF-18 to be required for the age-dependent increase in the expression of representative detoxification enzymes and immune effectors. Accordingly, our functional analysis revealed a role for DAF-18 in the innate immunity of adult *C. elegans*. While DAF-18 had no bearing on the ability of larval stage animals to resist bacterial pathogens, postreproductive adults lacking functional DAF-18 died from *P. aeruginosa* infection more rapidly than their wildtype counterparts. This same age-dependent enhanced pathogen susceptibility is a characteristic of animals lacking functional DAF-16 (this study and McHugh *et al*. 2020). Our results therefore support the possibility that DAF-18 is necessary to activate DAF-16 during aging and that it contributes to DAF-16-mediated host defense in adult *C. elegans*.

PTEN is tightly regulated in mammals, yet comparatively few DAF-18 regulators have been identified in *C. elegans*. Since DAF-18 has not been extensively characterized during aging, our studies gave us an opportunity to search for possible mechanisms of DAF-18 regulation that had not previously been reported in worms. The results of our reverse genetic candidate gene approach suggest that in order to perform its functions in innate immunity during adulthood, DAF-18 must be deubiquitinated by MATH-33 and it must by sumoylated through conjugation to SMO-1. Based on their effects on PTEN, both of these modifications may be necessary to efficiently target DAF-18 to the plasma membrane. Monoubiquitination of PTEN promotes its translocation to the nucleus, and polyubiquitinated PTEN is degraded (Chen *et al*. 2018). Therefore, regardless of which ubiquitin moiety is removed from the protein, the result is to promote its activity in the cytosol. Binding of PTEN to the plasma membrane is associated with its sumoylation on two adjacent lysine residues (Huang *et al*. 2012). If the effects of these posttranslational modifications are evolutionarily conserved, de-ubiquitinated, sumoylated DAF-18 should be closely positioned to its lipid substrate. This provides additional evidence to suggest that DAF-18 functions as a lipid phosphatase during aging. One caveat of our data is that the knockdown of *math-33* and *smo-1* likely affects posttranslational modifications of multiple proteins and not only DAF-18. A notable case in point is that DAF-16 is a known substrate of MATH-33 under conditions of low IIS (Heimbucher *et al*. 2015). Additional studies are needed to confirm that the enhanced pathogen susceptibility phenotype caused by knockdown of *smo-1* and *math-33* in Day 6 adults is attributable solely to aberrant modification of DAF-18 that prevents DAF-16 activation. Should this be confirmed, then MATH-33 and the SUMO conjugating enzyme UBC-9 will represent new components of the regulatory mechanism governing DAF-18 function during aging.

Although DAF-18 is capable of dephosphorylating protein substrates, its role in lifespan determination requires its lipid phosphatase activity (Solari *et al*. 2005; Brisbin *et al*. 2009; Liu *et al*. 2014). This strongly suggests that the mechanism by which DAF-18 regulates DAF-16 is by dephosphorylating PIP_3_ produced by AGE-1, the PI3 kinase that is immediately downstream of the DAF-2 insulin receptor, thereby preventing activation of the inhibitory AKT-1 kinase. The requirement for DAF-18 to activate DAF-16 during adulthood therefore implies that there is a repressive signal through IIS at that time that must be opposed by DAF-18. If this is the case, then manipulations to relieve such repression should cause hyperactivation of DAF-16 and thus bring about phenotypes associated with its constitutive activity, including increased stress resistance. Indeed, we found that RNAi knockdown of *age-1* caused Day 6 adult but not larval stage *C. elegans* to become more resistant to bacterial infection. Similarly, knocking down *daf-2* beginning at the L1 stage improves the resistance of Day 6 animals to *P. aeruginosa* infection (McHugh *et al*. 2020). When we probed the specific timing requirements for *age-1*, *daf-18*, and *daf-16* in innate immunity using short pulses of RNAi, we found that we could delay the knockdown of *daf-16* and *daf-18* until the fourth day of adulthood and still recapitulate the effect of longer RNAi treatments targeting those genes but that this was too late to bring about the *age-1* knockdown phenotype. This indicates that by Day 4 of adulthood, AGE-1 has already completed its regulatory function whereas DAF-18 and DAF-16 remain active. Therefore, inhibition through IIS may be restricted to the first few days of adulthood. Supporting this possibility, the latest point at which *daf-2* knockdown can be initiated and produce the same magnitude of lifespan extension as in *daf-2* mutants is Day 3 of adulthood (Dillin *et al*. 2002). By Day 8 of adulthood, RNAi targeting *daf-2* has no effect on longevity. Why, then, is DAF-18 required after upstream steps in the IIS pathway have apparently stopped functioning? The consensus of several genetic analyses conducted by different teams of investigators is that *daf-18* mutations in fact only partially suppress mutations in *daf-2* and *age-1* (Larsen *et al*. 1995; Ogg and Ruvkun 1998; Honda and Honda 2002; Patel *et al*. 2008). One explanation for this phenomenon is that in the absence of PIP_3_ produced by AGE-1, a parallel compensatory pathway may be induced to activate PDK-1 and this, too, must be counteracted by DAF-18 (Ogg and Ruvkun 1998). Perhaps this is the scenario at later stages of adulthood once the IIS-mediated inhibition of DAF-16 has dissipated.

A repressive signal that acts through the IIS pathway to inhibit DAF-16 would originate with the production of insulins that act as DAF-2 agonists. There are 40 genes in the *C. elegans* genome that encode insulin-like peptides (ILPs), and they can either stimulate or block IIS (ref). While some insulins are expressed in intestinal cells, the majority are expressed in neurons. The prevailing model for lifespan determination in *C. elegans* is that DAF-2 agonist insulins secreted by chemosensory neurons in the worm’s head bind to DAF-2 in target tissues, including the intestine, to activate the IIS pathway. As a result, AKT-1 adds an inhibitory phosphate to DAF-16, sequestering it in the cytosol. Without the robust expression of DNA repair machinery, chaperones, detoxification enzymes, immune effectors, and other DAF-16 transcriptional targets important for defending against stress or mitigating its effects, the capacity of animals to withstand challenges is hobbled, leaving them vulnerable to cellular damage that leads to functional decline. Our data indicate that while this may be the case very early in adulthood, inhibition of DAF-16 is swiftly reversed potentially through an analogous neuroendocrine route.

In Day 6 adult animals we detected DAF-18::GFP in head neurons and body wall muscle. Although we could not empirically identify the specific neurons in which DAF-18 is expressed, based on their morphology and previous characterizations of DAF18, they are almost certain to be chemosensory amphid neurons that detect either volatile or soluble environmental cues (Tomioka *et al*. 2006; Brisbin *et al*. 2009). Using tissue-specific RNAi, we confirmed that neuronal DAF-18 is indeed functional and that it contributes to innate immunity in adult *C. elegans*. Those studies also established the intestine as a site of action for DAF-18 during adulthood, even though we were unable to detect its expression there by microscopy. Significantly, our data indicate that DAF-16 functions in the same tissues as DAF-18 in Day 6 animals. The neurons and intestine are notable in the context of aging for two reasons. First, there is an axis of communication between neurons (the brain) and the intestine, as described above, that may control organismal aging. The so-called gut-brain axis (GBA) is evolutionarily conserved and is implicated in some age-related diseases (Houser and Tansey 2017). Second, the activity of DAF-16 in the intestine or the neurons alone is sufficient to affect lifespan. Specifically, genetic manipulations in the *daf-2* mutant background to activate or inactivate DAF-16 exclusively in either the neurons or the intestine but no other tissues partially recapitulates or partially suppresses the *daf-2* longevity phenotype, respectively (Wolkow *et al*. 2000; Libina *et al*. 2003). Our data suggest that DAF-18 and DAF-16 cooperate in chemosensory neurons and the intestine during adulthood. We therefore envision a revised model for the neuroendocrine control of lifespan in *C. elegans*. In adult animals, DAF-18 in the neurons may be required for DAF-16 to upregulate the expression of a DAF-2 antagonist. This would oppose inhibition through IIS in distal tissues, including the intestine, leading to the activation of DAF-16 in a cell non-autonomous manner. This is consistent with the results of tissue-specific rescue experiments demonstrating that DAF-18 in the neurons can promote DAF-16 nuclear localization in other tissues (Masse *et al*. 2005). Within the intestine, DAF-18 may be required to reduce PIP_3_ levels produced by AGE-1 or by other compensatory pathways, permitting DAF-16 to upregulate the genes that preserve homeodynamic space and thus assure normal lifespan.

Proto-oncogenes are typically regarded as one of the best examples of such genes, as many are necessary for cell cycle progression earlier in life, yet their activity in older animals can lead to tumor progression. Tumor suppressors, on the other hand, play the opposite role. During development these genes prevent overgrowth of cells and limit organ size, and their absence causes developmental abnormalities or embryonic lethality (Pomerantz and Blau 2013). True to their name, later in life tumor suppressors stop aberrant cell division, thus helping to prevent cancer. Unlike proto-oncogenes, whose inactivation during aging would promote longevity by reducing the chances of developing life-threatening disease, organisms benefit from tumor suppressors being active both early and later in life. Our work on DAF-18, the *C. elegans* ortholog of the human tumor suppressor PTEN, demonstrates that this is the case even in animals that do not maintain mitotic cell populations during adulthood. While DAF-18 is important for the development of several tissues and controls cell proliferation in the germline, we find that it is required specifically during adulthood to protect animals from bacterial infection that would otherwise shorten their lifespan. This suggests that there is an evolutionarily conserved requirement for tumor suppressors to preserve lifespan, not just to prevent diseases of aging, and that their functional repertoire may expand later in life.

## ACKNOWLEDGEMENTS

We thank Rebecca Rivard for comments on the manuscript and other members of the Youngman lab for helpful discussions. KC was funded through the Beckman Scholars Program of the Arnold and Mabel Beckman Foundation. We acknowledge the office of the Dean of the College of Liberal Arts and Sciences at Villanova for additional funding. Some strains were provided by the CGC, which is funded by NIH Office of Research Infrastructure Programs (P40 OD010440). We are grateful to Alex Hajnal’s laboratory for sharing the DAF-18::GFP reporter strain with us.

---

**Figure S1.**
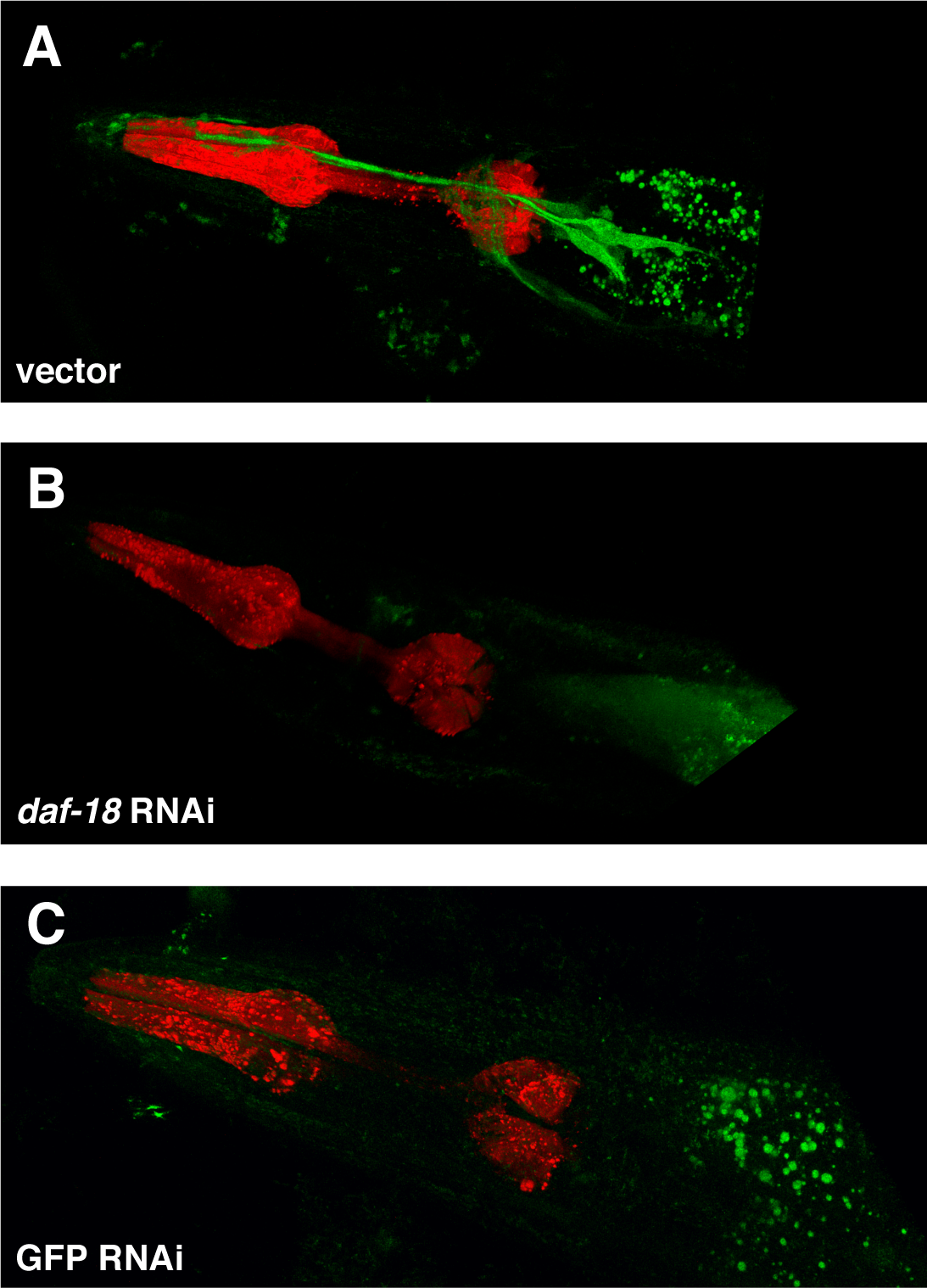
GFP expression in the head neurons of transgenic adult *C. elegans* is specific to DAF-18::GFP. *C. elegans* expressing the DAF-18::GFP translational fusion were subjected to feeding-based RNAi beginning at the L1 larval stage. At Day 6 of adulthood, animals treated with an empty vector (A) or with RNAi targeting either *daf-18* (B) or *gfp* (C) were imaged by confocal microscopy. Images are representative of animals examined during three independent biological replicates.

**Figure S2.**
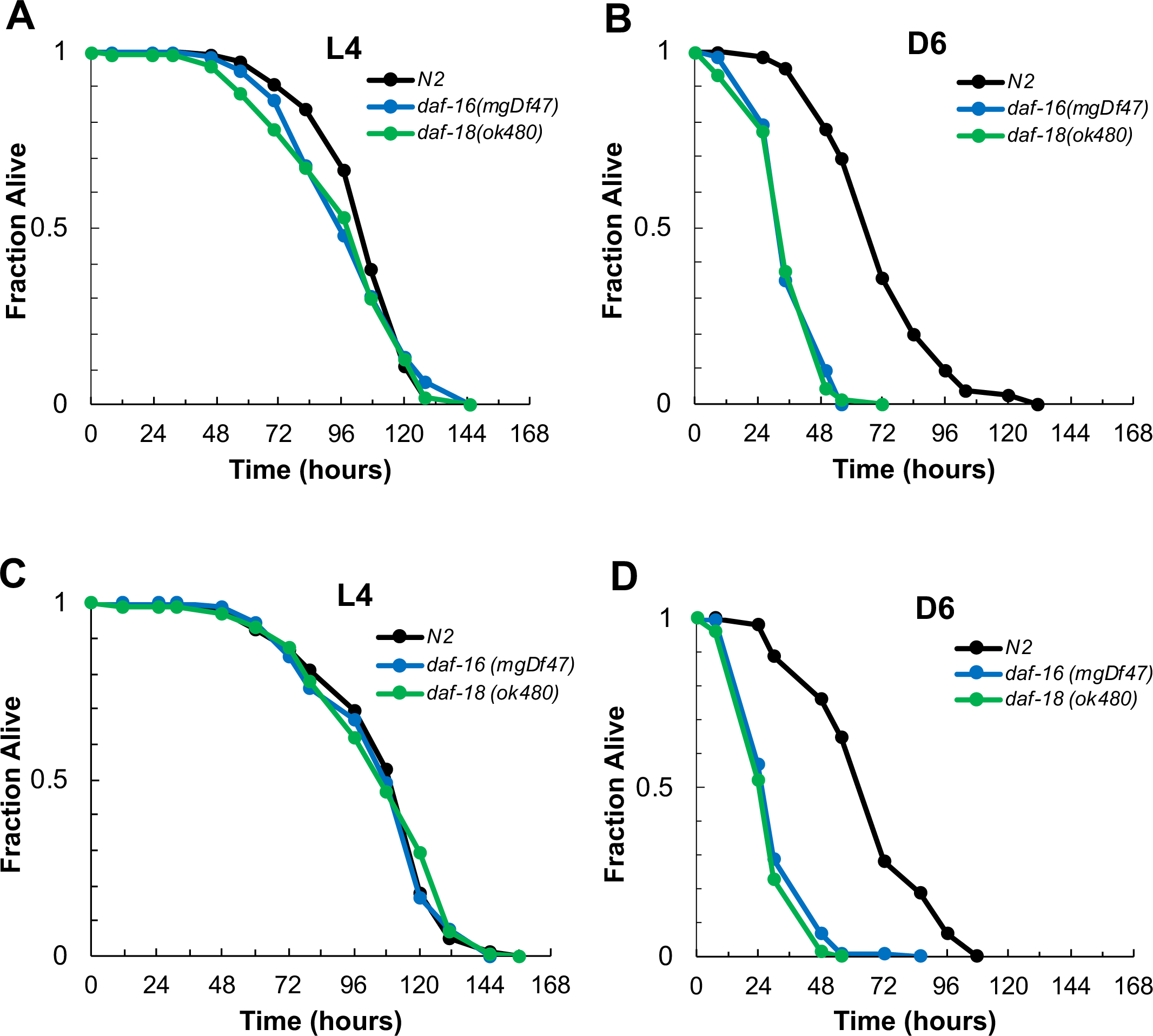
Additional replicates of *Pseudomonas aeruginosa* infection of larval and adult *daf-18* mutants. The survival of *daf-18(ok480)* mutants (blue) was compared to that of *daf-16(mgDf47)* animals (green) and to wildtype N2 animals *(black)* after infection with *P. aeruginosa* at either the L4 stage (A and C) or at Day 6 of adulthood (B and D). The fraction of animals alive is plotted as a function of time. Data from two independent biological replicates (A and B; C and D) are shown.

**Figure S3.**
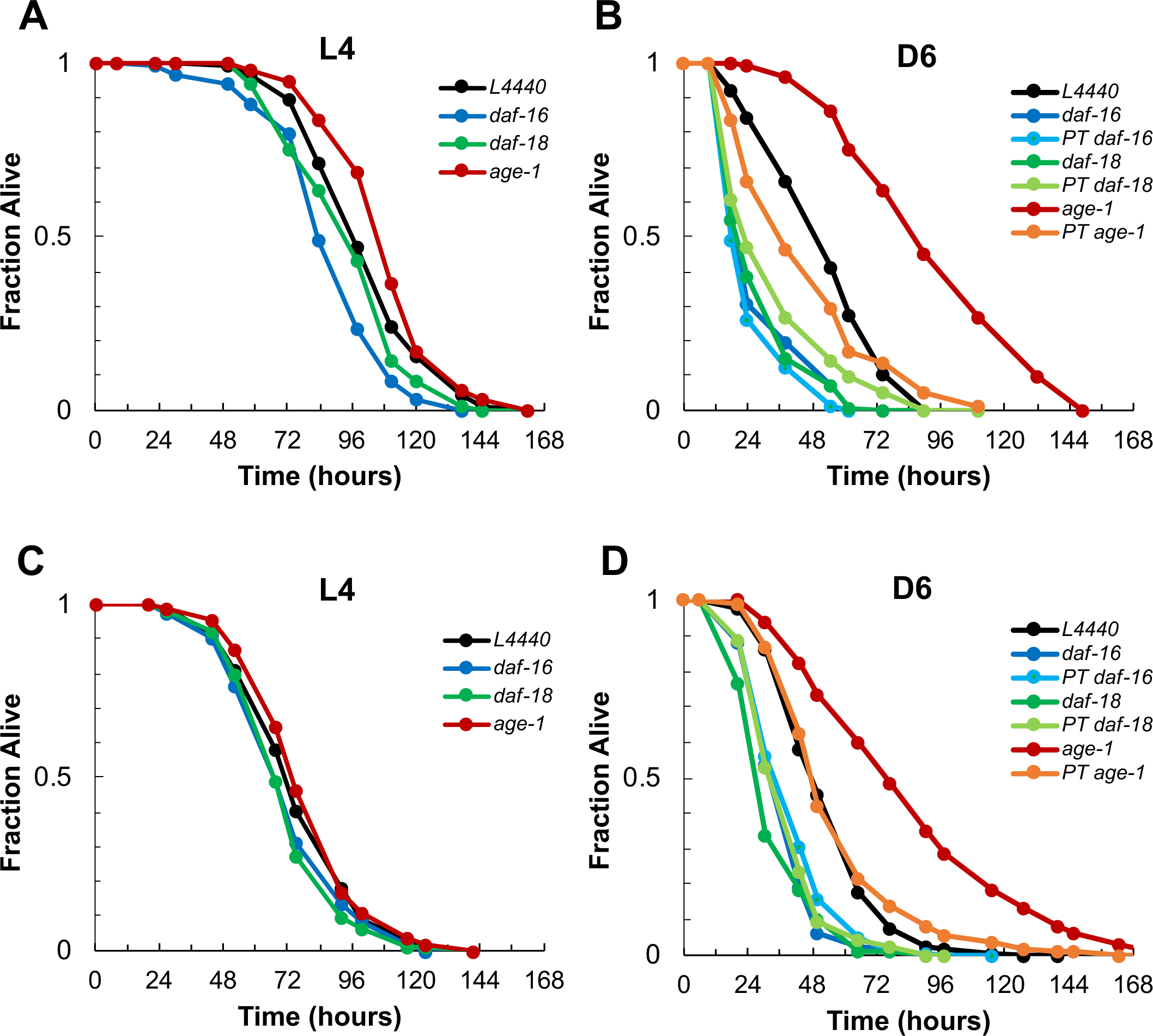
Additional replicates of RNAi experiment to investigate the timing requirement for *daf-18* in *C. elegans* innate immunity. Beginning at the L1 larval stage N2 wildtype animals were treated with an empty RNAi vector (L4440, black) or with RNAi targeting *daf-16* (dark blue), *daf-18* (dark green), or *age-1* (red) and then infected with *P. aeruginosa* at either the L4 larval stage (A, C) or at Day 6 of adulthood (B, D). Another group of animals was allowed to age on their normal *E. coli* food source until Day 4 of adulthood when RNAi targeting *daf-16* (light blue), *daf-18* (light green), or *age-1* (orange) was initiated. These animals were then infected with *P. aeruginosa* at Day 6 and their survival was monitored over time (B, D). Plotted in each panel is the fraction of animals alive as a function of time after the start of the infection. Data from two independent biological replicates (A and B; C and D) are shown.

**Figure S4.**
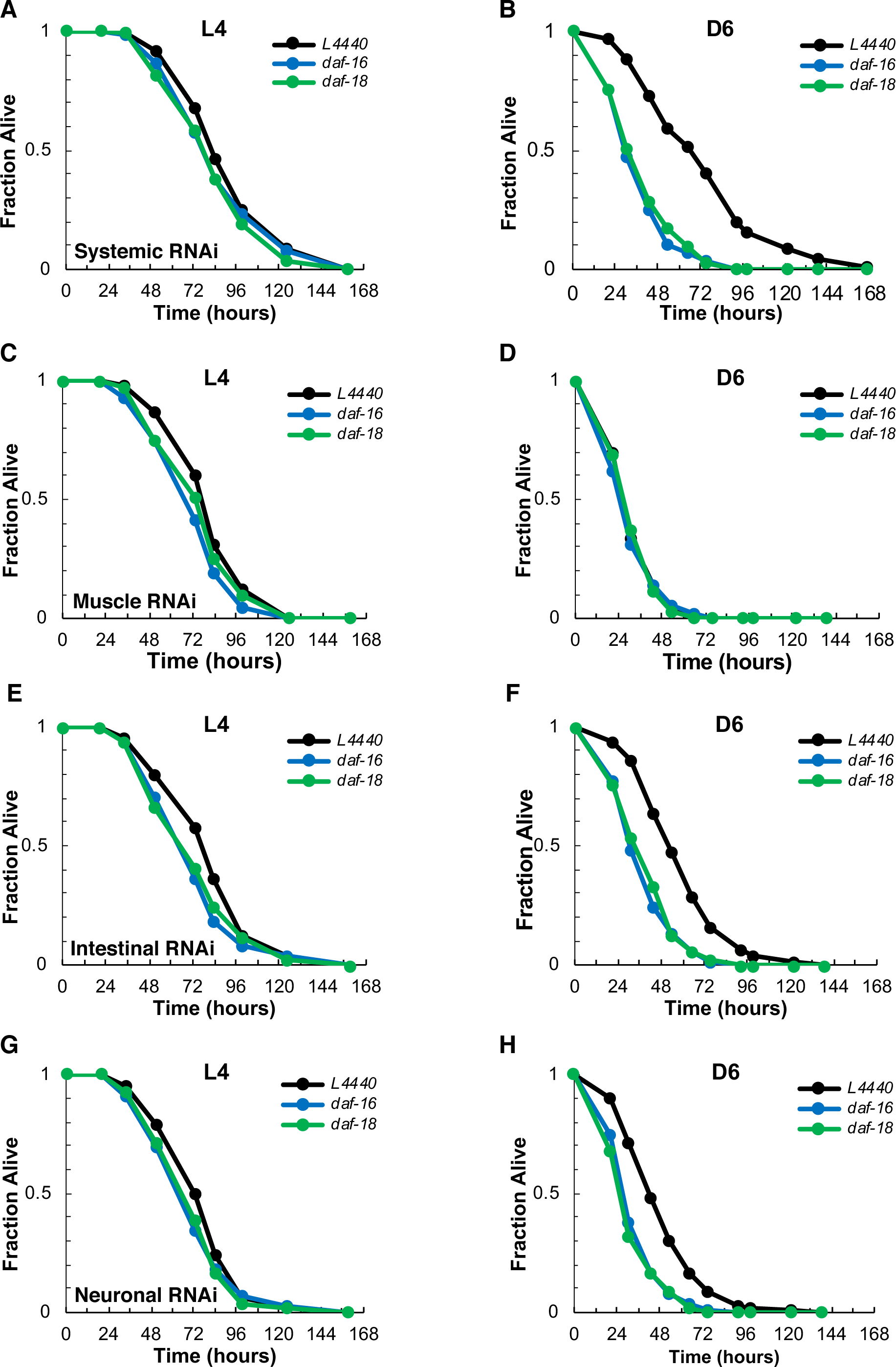
Second biological replicate of tissue-specific RNAi experiment to determine the site of action of *daf-18* in innate immunity during adulthood. N2 wild type animals (A and B) along with mutants in which RNAi-mediated gene knockdowns are restricted to the muscle (C and D), the intestine (E and F) or the neurons (G and H) were treated with an empty RNAi vector (L4440, black) or with constructs targeting *daf-18* (green) or *daf-16* (blue) starting at the L1 larval stage. *P. aeruginosa* infection was initiated at either the L4 larval stage (A, C, E, G) or at Day 6 of adulthood (B, D, F, H) and the survival of infected animals was recorded over time. In each panel, the fraction of worms alive is plotted as a function of time after the start of the infection.

**Figure S5.**
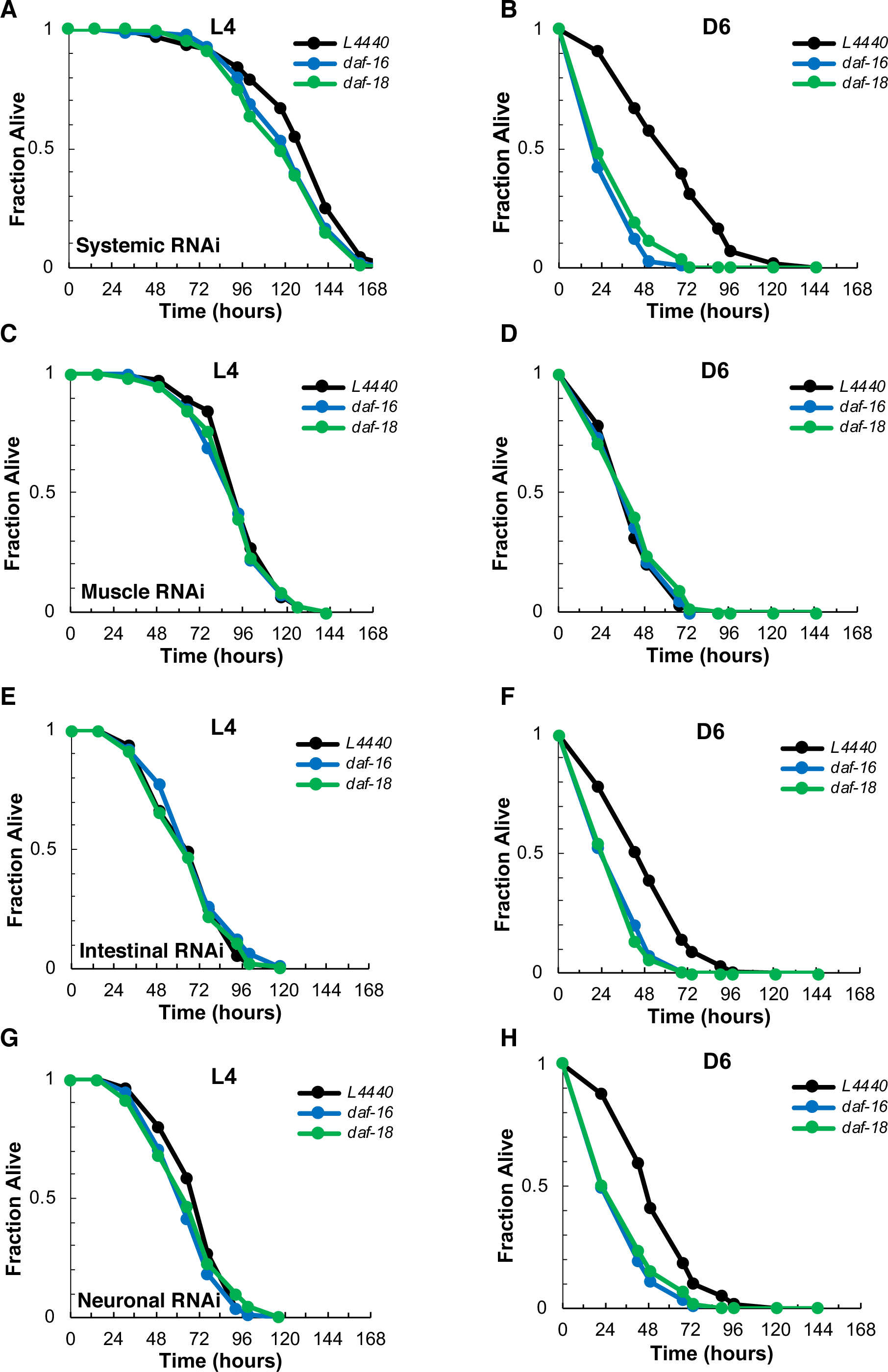
Third biological replicate of tissue-specific RNAi experiment to determine the site of action of *daf-18* in innate immunity during adulthood. N2 wild type animals (A and B) along with mutants in which RNAi-mediated gene knockdowns are restricted to the muscle (C and D), the intestine (E and F) or the neurons (G and H) were treated with an empty RNAi vector (L4440, black) or with constructs targeting *daf-18* (green) or *daf-16* (blue) starting at the L1 larval stage. *P. aeruginosa* infection was initiated at either the L4 larval stage (A, C, E, G) or at Day 6 of adulthood (B, D, F, H) and the survival of infected animals was recorded over time. In each panel, the fraction of worms alive is plotted as a function of time after the start of the infection.

**Figure S6.**
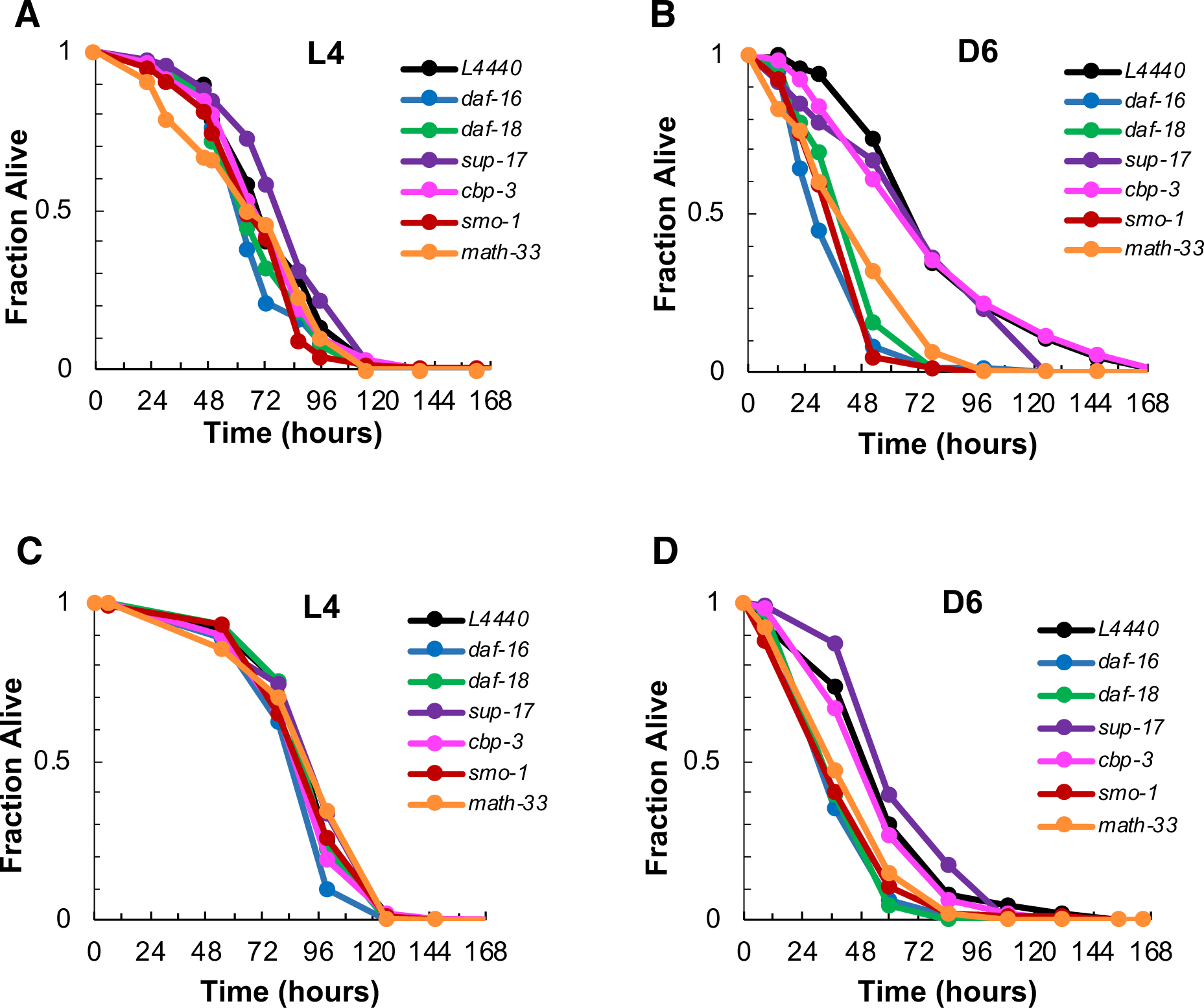
**Additional replicates of RNAi experiment to functionally characterize putative regulators of DAF-18.** RNAi treatment to knock down the candidate DAF-18 regulators *sup-17* (purple), *cbp-3* (dark green), *smo-1* (red), or *math-33* (yellow) was initiated in N2 wildtype *C. elegans* at the L1 larval stage. For comparison, other animals were treated with an empty vector (L4440, black) or with RNAi constructs targeting *daf-18* (light green) or *daf-16* (blue). Animals were then infected with *P. aeruginosa* as either L4 larvae (A, C) or as Day 6 adults (B, D), and the fraction of worms alive was recorded at regular intervals until all individuals had died. Data from two independent biological replicates (A and B; C and D) are shown.

**Table S1.** Tabulation of scoring data from manual assessment of *plys-7::GFP* expression shown in Fig. 2. The numbers of animals assigned to each category of *plys-7::GFP* expression (low, medium, or high) are reported according to age and RNAi treatement for three independent biological replicates. Separate tables show the total number of worms in each category across all three replicates and the proportions of animals in each category (labled as “converted to fractions”).

Larval Stage 4
|  | Low | Med | High |
| --- | --- | --- | --- |
| L4440 | 194 | 6 | 0 |
| daf-16 | 195 | 5 | 0 |
| daf-18 | 195 | 4 | 1 |
| age-1 | 197 | 3 | 0 |

Day 3 of Adulthood
|  | Low | Med | High |
| --- | --- | --- | --- |
| L4440 | 9 | 88 | 103 |
| daf-16 | 153 | 44 | 3 |
| daf-18 | 126 | 72 | 2 |
| age-1 | 0 | 10 | 190 |

Day 6 of Adulthood
|  | Low | Med | High |
| --- | --- | --- | --- |
| L4440 | 0 | 13 | 187 |
| daf-16 | 91 | 99 | 10 |
| daf-18 | 90 | 104 | 6 |
| age-1 | 0 | 11 | 189 |

Replicate 2 Larval Stage 4
|  | Low | Med | High |
| --- | --- | --- | --- |
| L4440 | 183 | 17 | 0 |
| daf-16 | 192 | 8 | 0 |
| daf-18 | 185 | 15 | 0 |
| age-1 | 198 | 2 | 0 |

Day 3 of Adulthood
|  | Low | Med | High |
| --- | --- | --- | --- |
| L4440 | 31 | 75 | 94 |
| daf-16 | 177 | 23 | 0 |
| daf-18 | 168 | 32 | 0 |
| age-1 | 5 | 17 | 178 |

Day 6 of Adulthood
|  | Low | Med | High |
| --- | --- | --- | --- |
| L4440 | 0 | 14 | 186 |
| daf-16 | 151 | 47 | 2 |
| daf-18 | 113 | 83 | 4 |
| age-1 | 0 | 4 | 196 |

Replicate 3 Larval Stage 4
|  | Low | Med | High |
| --- | --- | --- | --- |
| L4440 | 195 | 5 | 0 |
| daf-16 | 189 | 10 | 1 |
| daf-18 | 191 | 9 | 0 |
| age-1 | 190 | 9 | 1 |

Day 3 of Adulthood
|  | Low | Med | High |
| --- | --- | --- | --- |
| L4440 | 21 | 107 | 72 |
| daf-16 | 159 | 40 | 1 |
| daf-18 | 164 | 34 | 2 |
| age-1 | 2 | 12 | 186 |

Day 6 of Adulthood
|  | Low | Med | High |
| --- | --- | --- | --- |
| L4440 | 3 | 31 | 164 |
| daf-16 | 138 | 53 | 9 |
| daf-18 | 131 | 56 | 13 |
| age-1 | 0 | 5 | 195 |

Totals from all three replicates Larval Stage 4
|  | Low | Med | High |
| --- | --- | --- | --- |
| L4440 | 572 | 28 | 0 |
| daf-16 | 576 | 23 | 1 |
| daf-18 | 571 | 28 | 1 |
| age-1 | 585 | 14 | 1 |

Day 3 of Adulthood
|  | Low | Med | High |
| --- | --- | --- | --- |
| L4440 | 61 | 270 | 269 |
| daf-16 | 489 | 107 | 4 |
| daf-18 | 458 | 138 | 4 |
| age-1 | 7 | 39 | 554 |

Day 6 of Adulthood
|  | Low | Med | High |
| --- | --- | --- | --- |
| L4440 | 3 | 58 | 537 |
| daf-16 | 380 | 199 | 21 |
| daf-18 | 334 | 243 | 23 |
| age-1 | 0 | 20 | 580 |

Standard Deviations: Larval Stage 4
|  | Low | Med | High |
| --- | --- | --- | --- |
| L4440 | 6.658328 | 6.658328 | 0 |
| daf-16 | 3 | 2.516611 | 0.57735 |
| daf-18 | 5.033223 | 5.507571 | 0.57735 |
| age-1 | 4.358899 | 3.785939 | 0.57735 |

Day 3 of Adulthood
|  | Low | Med | High |
| --- | --- | --- | --- |
| L4440 | 11.01514 | 16.09348 | 15.94783 |
| daf-16 | 12.49 | 11.15049 | 1.527525 |
| daf-18 | 23.18045 | 22.53886 | 1.154701 |
| age-1 | 2.516611 | 3.605551 | 6.110101 |

Day 6 of Adulthood
|  | Low | Med | High |
| --- | --- | --- | --- |
| L4440 | 1.732051 | 10.11599 | 13 |
| daf-16 | 31.56475 | 28.44878 | 4.358899 |
| daf-18 | 20.55075 | 24.06242 | 4.725816 |
| age-1 | 0 | 3.785939 | 3.785939 |

Converted to Fractions\*\* Larval Stage 4
|  | Low | Med | High |
| --- | --- | --- | --- |
| L4440 | 0.97 | 0.03 | 0 |
| daf-16 | 0.975 | 0.025 | 0 |
| daf-18 | 0.975 | 0.02 | 0.005 |
| age-1 | 0.985 | 0.015 | 0 |

Day 3 of Adulthood
|  | Low | Med | High |
| --- | --- | --- | --- |
| L4440 | 0.045 | 0.44 | 0.515 |
| daf-16 | 0.765 | 0.22 | 0.015 |
| daf-18 | 0.63 | 0.36 | 0.01 |
| age-1 | 0 | 0.05 | 0.95 |

Day 6 of Adulthood
|  | Low | Med | High |
| --- | --- | --- | --- |
| L4440 | 0 | 0.065 | 0.935 |
| daf-16 | 0.455 | 0.495 | 0.05 |
| daf-18 | 0.45 | 0.52 | 0.03 |
| age-1 | 0 | 0.055 | 0.945 |

Larval Stage 4
|  | Low | Med | High |
| --- | --- | --- | --- |
| L4440 | 0.915 | 0.085 | 0 |
| daf-16 | 0.96 | 0.04 | 0 |
| daf-18 | 0.925 | 0.075 | 0 |
| age-1 | 0.99 | 0.01 | 0 |

Day 3 of Adulthood
|  | Low | Med | High |
| --- | --- | --- | --- |
| L4440 | 0.155 | 0.375 | 0.47 |
| daf-16 | 0.885 | 0.115 | 0 |
| daf-18 | 0.84 | 0.16 | 0 |
| age-1 | 0.025 | 0.085 | 0.89 |

Day 6 of Adulthood
|  | Low | Med | High |
| --- | --- | --- | --- |
| L4440 | 0 | 0.07 | 0.93 |
| daf-16 | 0.755 | 0.235 | 0.01 |
| daf-18 | 0.565 | 0.415 | 0.02 |
| age-1 | 0 | 0.02 | 0.98 |

Larval Stage 4
|  | Low | Med | High |
| --- | --- | --- | --- |
| L4440 | 0.975 | 0.025 | 0 |
| daf-16 | 0.945 | 0.05 | 0.005 |
| daf-18 | 0.955 | 0.045 | 0 |
| age-1 | 0.95 | 0.045 | 0.005 |

Day 3 of Adulthood
|  | Low | Med | High |
| --- | --- | --- | --- |
| L4440 | 0.105 | 0.535 | 0.36 |
| daf-16 | 0.795 | 0.2 | 0.005 |
| daf-18 | 0.82 | 0.17 | 0.01 |
| age-1 | 0.01 | 0.06 | 0.93 |

Day 6 of Adulthood
|  | Low | Med | High |
| --- | --- | --- | --- |
| L4440 | 0.015 | 0.155 | 0.82 |
| daf-16 | 0.69 | 0.265 | 0.045 |
| daf-18 | 0.655 | 0.28 | 0.065 |
| age-1 | 0 | 0.025 | 0.975 |

Averages: Larval Stage 4
|  | Low | Med | High |
| --- | --- | --- | --- |
| L4440 | 0.953333 | 0.046667 | 0 |
| daf-16 | 0.96 | 0.038333 | 0.001667 |
| daf-18 | 0.951667 | 0.046667 | 0.001667 |
| age-1 | 0.975 | 0.023333 | 0.001667 |

Larval Stage 4
|  | Low | Med | High |
| --- | --- | --- | --- |
| L4440 | 0.033292 | 0.033292 | 0 |
| daf-16 | 0.015 | 0.012583 | 0.002887 |
| daf-18 | 0.025166 | 0.027538 | 0.002887 |
| age-1 | 0.021794 | 0.01893 | 0.002887 |

Day 3 of Adulthood
|  | Low | Med | High |  | Low | Med | High |
| --- | --- | --- | --- | --- | --- | --- | --- |
| L4440 | 0.101667 | 0.45 | 0.448333 | L4440 | 0.055076 | 0.080467 | 0.079739 |
| daf-16 | 0.815 | 0.178333 | 0.006667 | daf-16 | 0.06245 | 0.055752 | 0.007638 |
| daf-18 | 0.763333 | 0.23 | 0.006667 | daf-18 | 0.115902 | 0.112694 | 0.005774 |
| age-1 | 0.011667 | 0.065 | 0.923333 | age-1 | 0.012583 | 0.018028 | 0.030551 |
| Day 6 of Adulthood |  |  |  | Day 6 of Adulthood |  |  |  |
|  | Low | Med | High |  | Low | Med | High |
| L4440 | 0.005 | 0.096667 | 0.895 | L4440 | 0.00866 | 0.05058 | 0.065 |
| daf-16 | 0.633333 | 0.331667 | 0.035 | daf-16 | 0.157824 | 0.142244 | 0.021794 |
| daf-18 | 0.556667 | 0.405 | 0.038333 | daf-18 | 0.102754 | 0.120312 | 0.023629 |
| age-1 | 0 | 0.033333 | 0.966667 | age-1 | 0 | 0.01893 | 0.01893 |

**Table S4.** Compiled fold differences in expression of mtl-1, sod-3, lys-7, and cpr-2 between L4 and Day 6 in animals treated with RNAi targeting the indicated genes for three independent biological replicates. These data correspond to the bar graph in Fig. 2E. Average fold difference in expression betweeh Day 6 and L4 with standard deviations and standard error are shown below.

| *mtl-1* | L4440 | *daf-16* | *daf-18* | *age-1* |
| --- | --- | --- | --- | --- |
| Rep 1 | 11.3137085 | 0.30707579 | 0.47521974 | 79.524822 |
| Rep 2 | 4.60542192 | 0.32458465 | 0.37892914 | 29.8570557 |
| Rep 3 | 5.00487256 | 0.29185695 | 0.44647976 | 69.8731689 |
| Av | 6.97466766 | 0.30783913 | 0.43354288 | 59.7516822 |
| st dev | 3.76302362 | 0.0163772 | 0.04943169 | 26.3354347 |

| *sod-3* | L4440 | *daf-16* | *daf-18* | *age-1* |
| --- | --- | --- | --- | --- |
| Rep 1 | 1.81923679 | 0.0900373 | 0.14899474 | 218.779437 |
| Rep 2 | 2.17849731 | 0.29456678 | 0.37371231 | 144.340599 |
| Rep 3 | 1.49484925 | 0.20073072 | 0.21563551 | 219.285509 |
| Av | 1.83086112 | 0.1951116 | 0.24611419 | 194.135181 |
| st dev | 0.34197224 | 0.10238046 | 0.11541754 | 43.124116 |

| *lys-7* | L4440 | *daf-16* | *daf-18* | *age-1* |
| --- | --- | --- | --- | --- |
| Rep 1 | 128.296085 | 10.1026836 | 14.6213032 | 543.699868 |
| Rep 2 | 77.171701 | 8.39773347 | 12.1257325 | 705.906519 |
| Rep 3 | 121.937664 | 10.362738 | 9.42613711 | 639.145241 |
| Av | 109.13515 | 9.6210517 | 12.0577243 | 629.583876 |
| st dev | 27.8631284 | 1.0673742 | 2.59825067 | 81.5249298 |

| *cpr-2* | L4440 | *daf-16* | *daf-18* | *age-1* |
| --- | --- | --- | --- | --- |
| Rep 1 | 12.2950015 | 3.03844533 | 3.24900959 | 112.465082 |
| Rep 2 | 25.9320916 | 2.73208051 | 4.0278222 | 144.007486 |
| Rep 3 | 15.5984777 | 4.00925265 | 6.36429187 | 129.487288 |
| Av | 17.9418569 | 3.25992616 | 4.54704122 | 128.653285 |
| st dev | 7.11414972 | 0.66677019 | 1.62124558 | 15.7877321 |

| Averages | *mtl-1* | *sod-3* | *lys-7* | *cpr-2* |
| --- | --- | --- | --- | --- |
| L4440 | 6.97466766 | 1.83086112 | 109.13515 | 17.9418569 |
| *daf-16* | 0.30783913 | 0.1951116 | 9.6210517 | 3.25992616 |
| *daf-18* | 0.43354288 | 0.24611419 | 12.0577243 | 4.54704122 |
| *age-1* | 59.7516822 | 194.135181 | 629.583876 | 128.653285 |

| Standard Deviations | *mtl-1* | *sod-3* | *lys-7* | *cpr-2* |
| --- | --- | --- | --- | --- |
| L4440 | 3.76302362 | 0.34197224 | 27.8631284 | 7.11414972 |
| *daf-16* | 0.0163772 | 0.10238046 | 1.0673742 | 0.66677019 |
| *daf-18* | 0.04943169 | 0.11541754 | 2.59825067 | 1.62124558 |
| *age-1* | 26.3354347 | 43.124116 | 81.5249298 | 15.7877321 |

| Standard Err | *mtl-1* | *sod-3* | *lys-7* | *cpr-2* |
| --- | --- | --- | --- | --- |
| L4440 | 2.1725827 | 0.19743776 | 16.0867847 | 4.10735626 |
| *daf-16* | 0.00945538 | 0.05910938 | 0.61624878 | 0.38495995 |
| *daf-18* | 0.0285394 | 0.06663635 | 1.50010072 | 0.93602657 |
| *age-1* | 15.2047703 | 24.89772 | 47.0684402 | 9.11505139 |

**Table S5.**
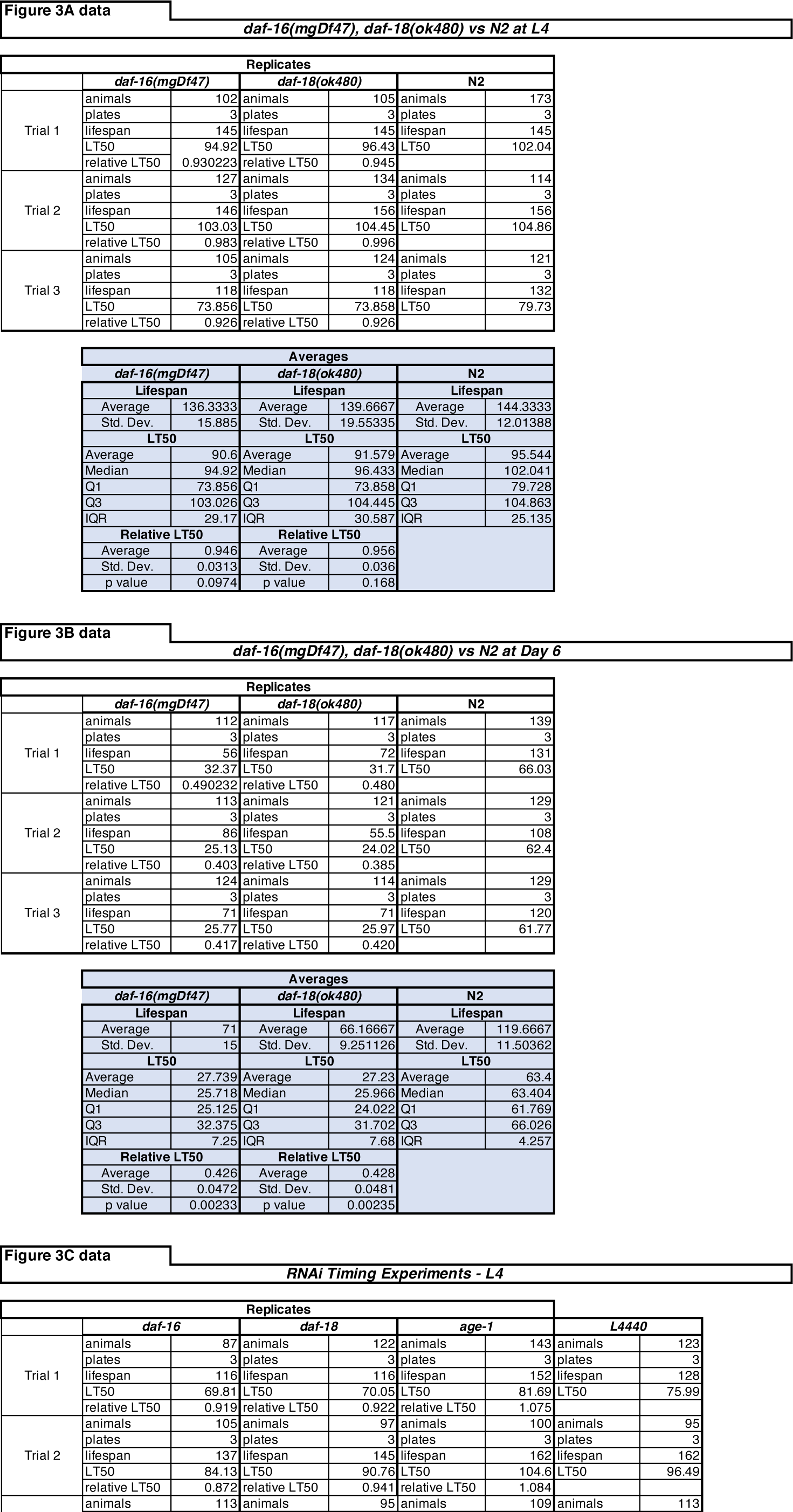
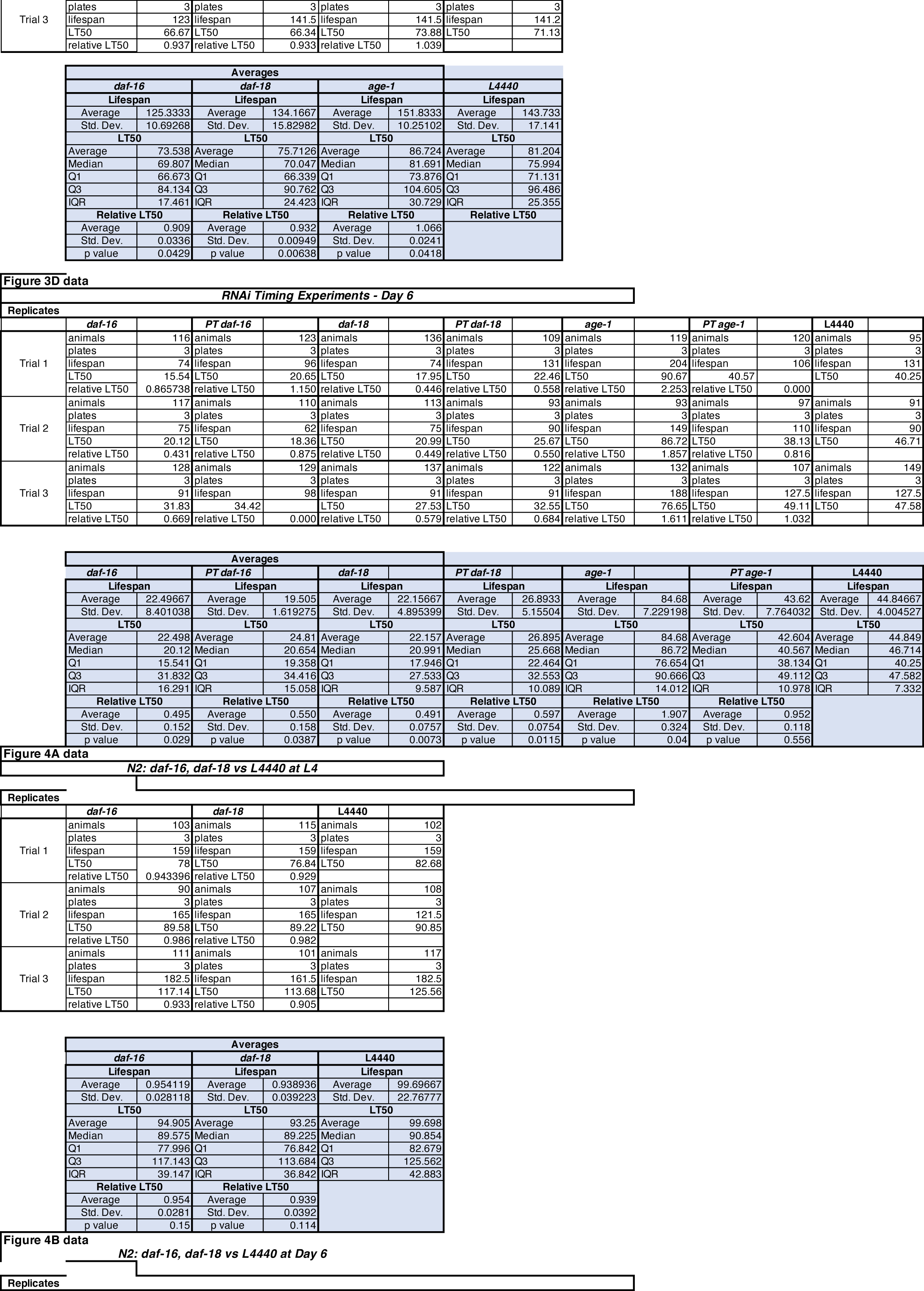
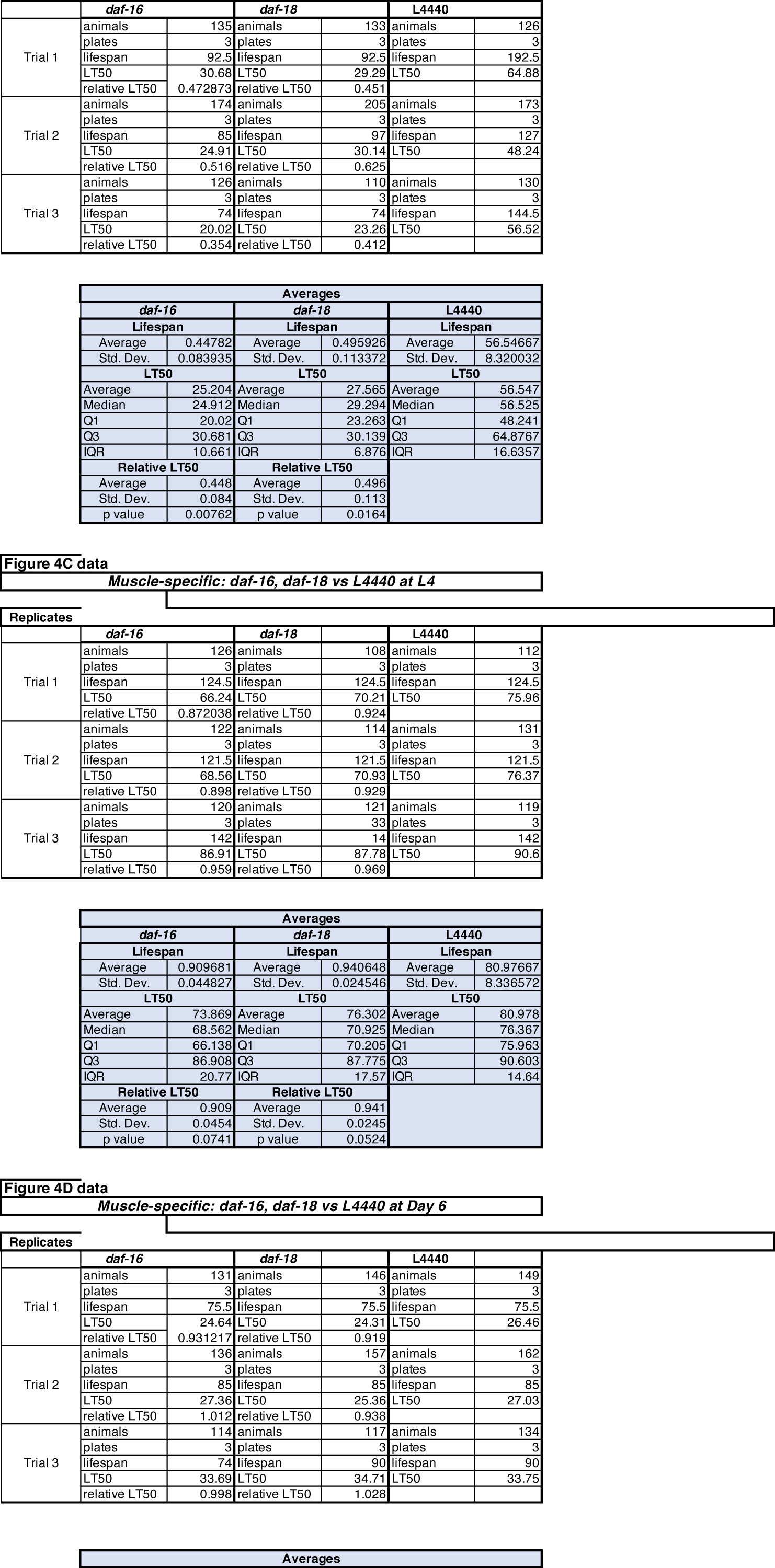
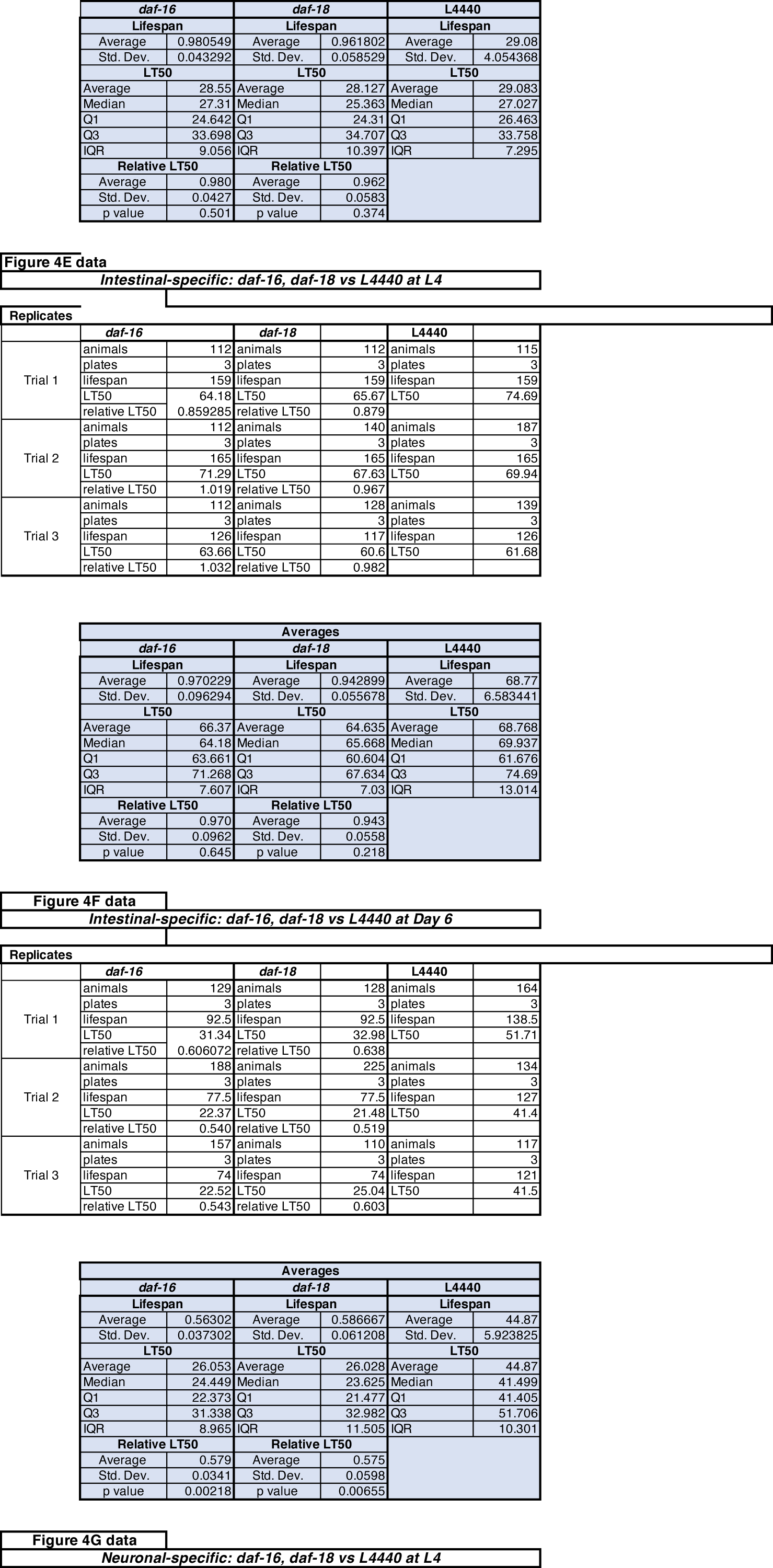
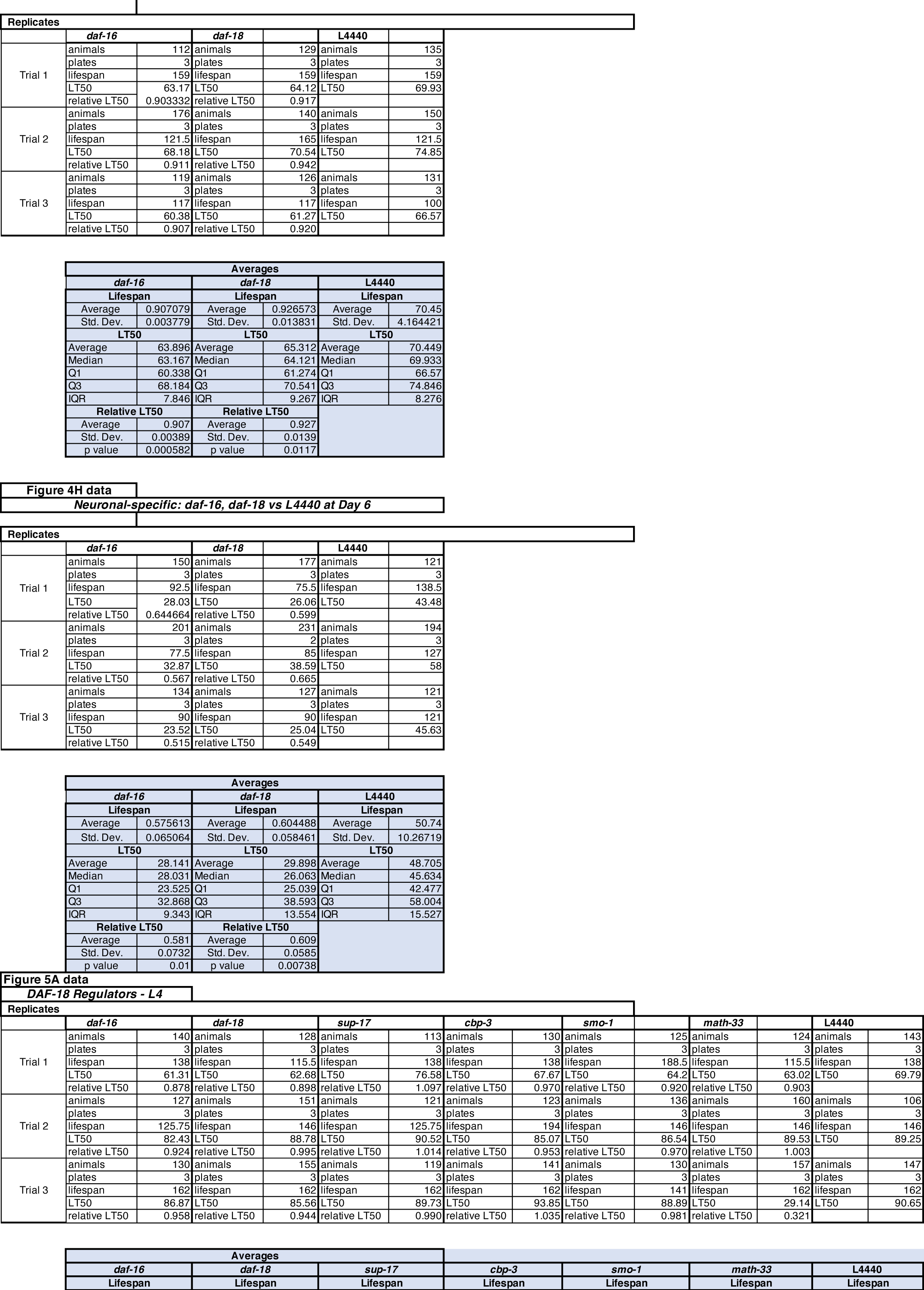
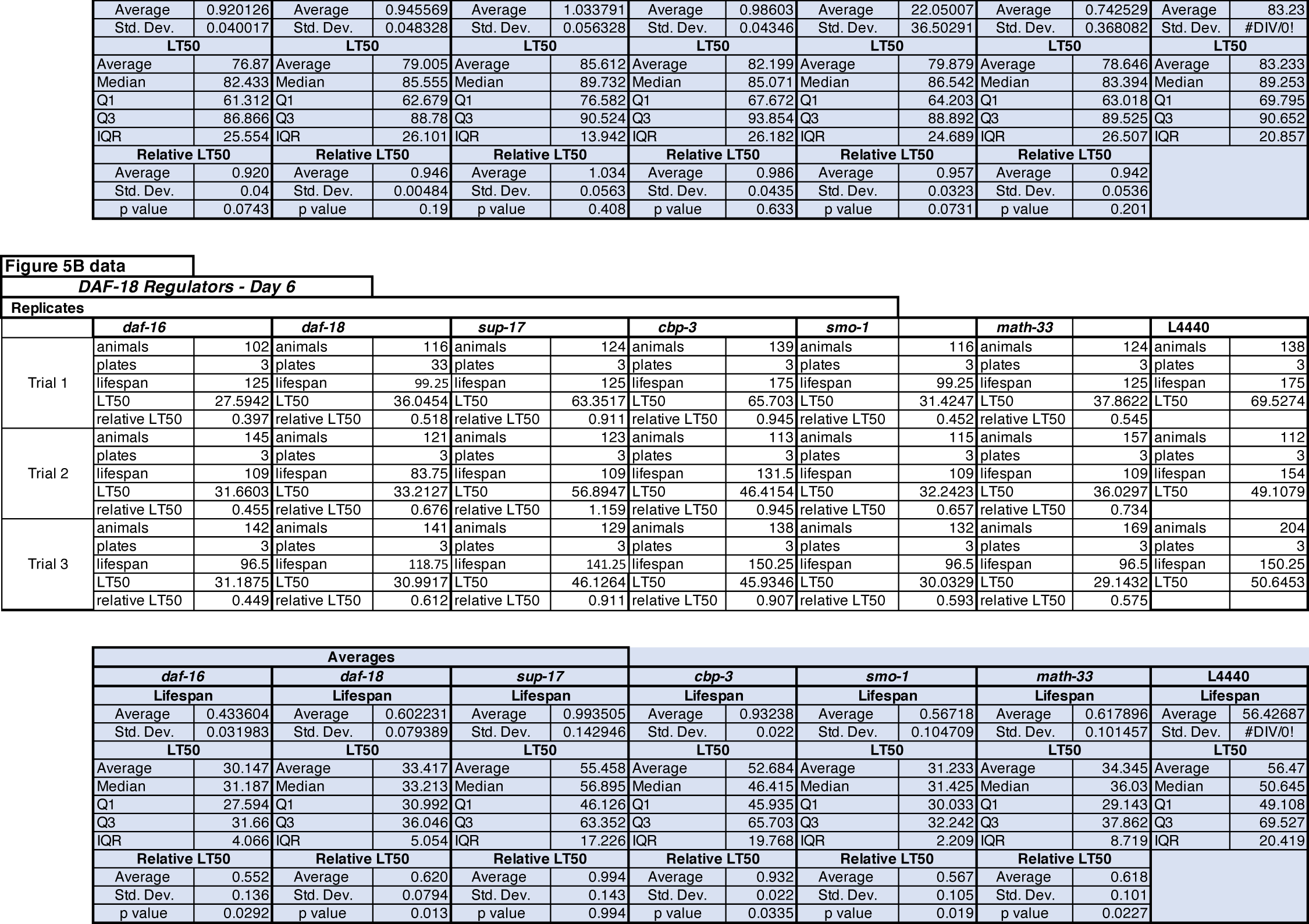
Experimental details and statistical analyses of *P. aeruginosa* infection assays corresponding to survival curves shown in Figs. 3-5. For each surival curve depicted in the figures, supporting information regarding the number of animals in each replicate, their maximum lifespan and their median lifespan (LT50) is provided. Relative LT50 was calculated by dividing the LT50 of an experimental group (mutant or RNAi-treated animals) by the LT50 of the corresponding control group. Compiled data for all replicates for a particular experiment are provided in the Averages section of each entry, which also includes statistical analyses. Lifespan and LT50 are presented in units of hours. p-values were determined using one-sample t-test. Std. dev., standard deviation; Q1, first quartile; Q3, third quartile; IQR, interquartile range

## REFERENCES

Chen, C.-Y., J. Chen, L. He, and B. L. Stiles, 2018 PTEN: Tumor Suppressor and Metabolic Regulator. Front. Endocrinol. 9:.

Dorman, J. B., B. Albinder, T. Shroyer, and C. Kenyon, 1995 The age-1 and daf-2 genes function in a common pathway to control the lifespan of Caenorhabditis elegans. Genetics 141: 1399–1406.

Huang, J., J. Yan, J. Zhang, S. Zhu, Y. Wang et al., 2012 SUMO1 modification of PTEN regulates tumorigenesis by controlling its association with the plasma membrane. Nat Commun 3: 1–12.

Libina, N., J. R. Berman, and C. Kenyon, 2003 Tissue-specific activities of C. elegans DAF-16 in the regulation of lifespan. Cell 115: 489–502.

McHugh, D. R., E. Koumis, P. Jacob, J. Goldfarb, M. Schlaubitz-Garcia et al., 2020 DAF-16 and SMK-1 Contribute to Innate Immunity During Adulthood in Caenorhabditis elegans. G3: Genes, Genomes, Genetics.

Ogg, S., and G. Ruvkun, 1998 The C. elegans PTEN Homolog, DAF-18, Acts in the Insulin Receptor-like Metabolic Signaling Pathway. Molecular Cell 2: 887–893.

Wen, C., M. M. Metzstein, and I. Greenwald, 1997 SUP-17, a Caenorhabditis elegans ADAM protein related to Drosophila KUZBANIAN, and its role in LIN-12/NOTCH signalling. Development 124: 4759–4767.

Wolkow, C. A., K. D. Kimura, M.-S. Lee, and G. Ruvkun, 2000 Regulation of C. elegans Life-Span by Insulinlike Signaling in the Nervous System. Science 290: 147–150.

